# TIA1 Mediates Divergent Inflammatory Responses to Tauopathy in Microglia and Macrophages

**DOI:** 10.1101/2024.11.06.622325

**Authors:** Chelsea J. Webber, Sophie J. F. van de Spek, Anna Lourdes Cruz, Sambhavi Puri, Cheng Zhang, Jacqueline T. M. Aw, Georgia-Zeta Papadimitriou, Rebecca Roberts, Kiki Jiang, Thuc Nhan Tran, Lushuang Zhang, Alexandria Taylor, Zihan Wang, Jacob Porter, Ionnis Sotiropoulos, Andrew Emili, Joana Silva, Hu Li, Benjamin Wolozin

**Author notes:** Corresponding Author: Benjamin Wolozin, M.D., Ph.D., Professor, Dept. Of Anatomy & Neurobiology, Dept. of Neurology, Center for Systems Neuroscience, Neurophotonics Center Boston University Chobanian and Avedisian School of Medicine, 700 Albany St., W201 Boston, MA 02118-2526. Co-first authors (Chelsea J. Webber, Sophie J. F. van de Spek).

## Abstract

The RNA binding protein TIA1 is known to regulate stress responses. Here we show that TIA1 plays a much broader role in inflammatory cells, being required for the microglial sensome. We crossed TIA1 cKO mice (using a CX3CR1 driven cre element) to PS19 MAPT P301S tauopathy mice. The peripheral macrophages of TIA1 cKO mice exhibited a hyper-inflammatory phenotype with increased cytokine signaling, as expected. Surprisingly, the brains of these mice showed striking reductions in inflammation, including decreases in microglial inflammatory cytokines (TNFα and IL-1β) and sensome markers (CLEC7A, TREM2, ITGAX); these reductions were accompanied by corresponding decreases in tau pathology. Analysis of the brain TIA1 protein interactome identified brain selective TIA1 protein mediated pathways, including strong interactions with the microglial protein C1q, which directs pruning of dystrophic neurons. These results uncover a previously unknown regulatory role for TIA1 in microglial activation in the context of neurodegenerative disease and highlights the divergent regulation of two mononuclear phagocytic lineages: microglia and macrophages.

## Introduction

T-cell intracellular antigen-1 (TIA1) is an RNA-binding protein (RBP) that regulates splicing, transcription, and the translational stress response^1,2^. The pleiotropic actions of TIA1 obscure its roles in physiology and disease. TIA1 was originally identified as an RBP regulating the biology of the immune system^3^. Deletion of TIA1 rendered peripheral macrophages hyper-responsive to lipopolysaccharide (LPS) stimulation and, indeed, the TIA1 knockout mouse was proposed as a model of the autoimmune disease, rheumatoid arthritis^4^. TIA1 also plays an important role in promoting the translational stress response, where it functions as a core nucleating protein for membraneless granules termed stress granules^5^. The translational stress response plays an important role and is induced in response to the pathophysiology of neurodegenerative diseases^6,7^.

The importance of TIA1 in neurodegeneration is evident from studies of tau protein, which is one of the main pathological proteins contributing to Alzheimer’s disease (AD)^8–10^. Previously, we have shown that TIA1 co-localizes with tau pathology in the human brain^11^ and in animal models^12^, that recombinant tau and TIA1 are co-miscible in phase separated liquid droplets^13^, and that reducing TIA1 globally protects against disease progression in the P301S tau model of neurodegeneration^14^. However, the activity of TIA1 in both neurons and mononuclear phagocytic cells (including microglia and macrophages) limits interpretation of a global TIA1 KO. Microglia are the resident immune cells of the brain and are part of the macrophage lineage; microglia promote neuroinflammation and microgliosis during chronic stress^15–17^. Microglia become activated in response to protein aggregation and neurodegeneration, as well as diverse neuronal stress signals, including ATP/ADP release and activation of the complement cascade. The microglial genes whose transcripts increase in response to this activation have been categorized as “Disease Associated Microglial” (DAM) genes^18^. Chemokines and other stress signals attract microglia to the affected cells by activating receptors on microglia. These microglia-specific receptors are known as the microglial sensome, which partly overlaps with DAM genes^19,20^. In tauopathies, the microglial sensome detects tau aggregation and neuronal damage with receptors, such as triggering receptor expressed on myeloid cell 2 (TREM2)^21^. The sensome then promotes efferocytosis, inflammation and microgliosis. The microglial complement cascade also plays an important role in neurodegeneration, mediating synaptic pruning; C1q is correspondingly one of the proteins most consistently increased in neurodegenerative diseases^22^. Reducing chronic microglial activation and inflammation can delay neurodegeneration and reduces tau aggregation, although the effects of reducing efferocytosis vary depending on the nature of the pathology^23,24^.

We have shown that reducing TIA1 in neurons strongly protects against tau-mediated toxicity^14^. However, because TIA1 is a gene that is ubiquitously expressed, the specific role of TIA1 in microglia is unclear; does it inhibit immune responses as in peripheral macrophages or enhance stress responses as in neurons? To address this question, we created a mouse model that is a cross between the PS19 MAPT P301S tau mouse line and a mouse line with TIA1 selectively deleted in mononuclear phagocytic cells (i.e., microglia and macrophages) (P301S/TIA1cKO). Here we report that CX3CR1 driven knockout of TIA1 in microglia greatly reduces microglial inflammation, strongly inhibiting transcriptional networks including the C1q pathway. These results present with retention of a homeostatic morphology, reduction of transcripts for the microglial sensome, DAM genes cytokines and the complement response, as well as impacting direct protein interaction between microglial TIA1 and C1q. Analysis of the TIA1 brain protein interactome network identified C1q as the top interacting protein, as well as many synaptic TIA1 interacting proteins. Strikingly, the reduced microglial activation described above was accompanied by a decrease in tau pathology. The reduction of microglial responses observed with TIA1 cKO is accompanied by enhancement of peripheral macrophage activity in mice exposed to LPS, showing that TIA1 directs divergent phenotypic responses of inflammatory cells. These results highlight major differences in the regulation of microglia and peripheral macrophages, and point to TIA1 as a master regulator of monocyte phenotypes.

## Results

### Characterization of microglial TIA1 expression in TIA1cKO mice

TIA1 was conditionally knocked out in microglia using a *cre* recombinase under the control of the CX3CR1 regulatory sequences. Because CX3CR1 is expressed in myeloid-derived cells, the *cre* expression from the CX3CR1 promoter will also delete TIA1 in peripheral macrophages. P301S mice overexpressing human tau with a P301S mutation exhibit tau protein aggregation, neurodegeneration, cognitive dysfunction, and a have a shortened lifespan. To investigate the role of TIA1 in microglial function, P301S:TIA1^fl/fl^ mice were bred with TIA1^fl/fl^:CX3CR1^Cre/Cre^ mice to produce TIA1^fl/fl^:CX3CR1^-/Cre^ (TIA1cKO) and P301S:TIA1^fl/fl^:CX3CR1^-/Cre^ (P301S/TIA1cKO) mice. (**Figure 1A**). Microglia-selective TIA1 knockout in the brains of these mice was confirmed by first isolating microglia using CD11b+ magnetic beads and extracting RNA (**Figure 1B**). We measured the SIGLEC-H lectin marker to confirm that the CD11b+ fraction contained microglia (**Figure 1C**). TIA1 KO was confirmed in these microglia through rt-qPCR amplification of the TIA1 transcript (**Figure 1D**). There was no amplification of the TIA1 transcript indicating that TIA1 is depleted in the microglia of the TIA1cKO and P301S/TIA1cKO mice demonstrating successful removal of TIA1 from CX3CR1 positive cells.

**Figure 1.**
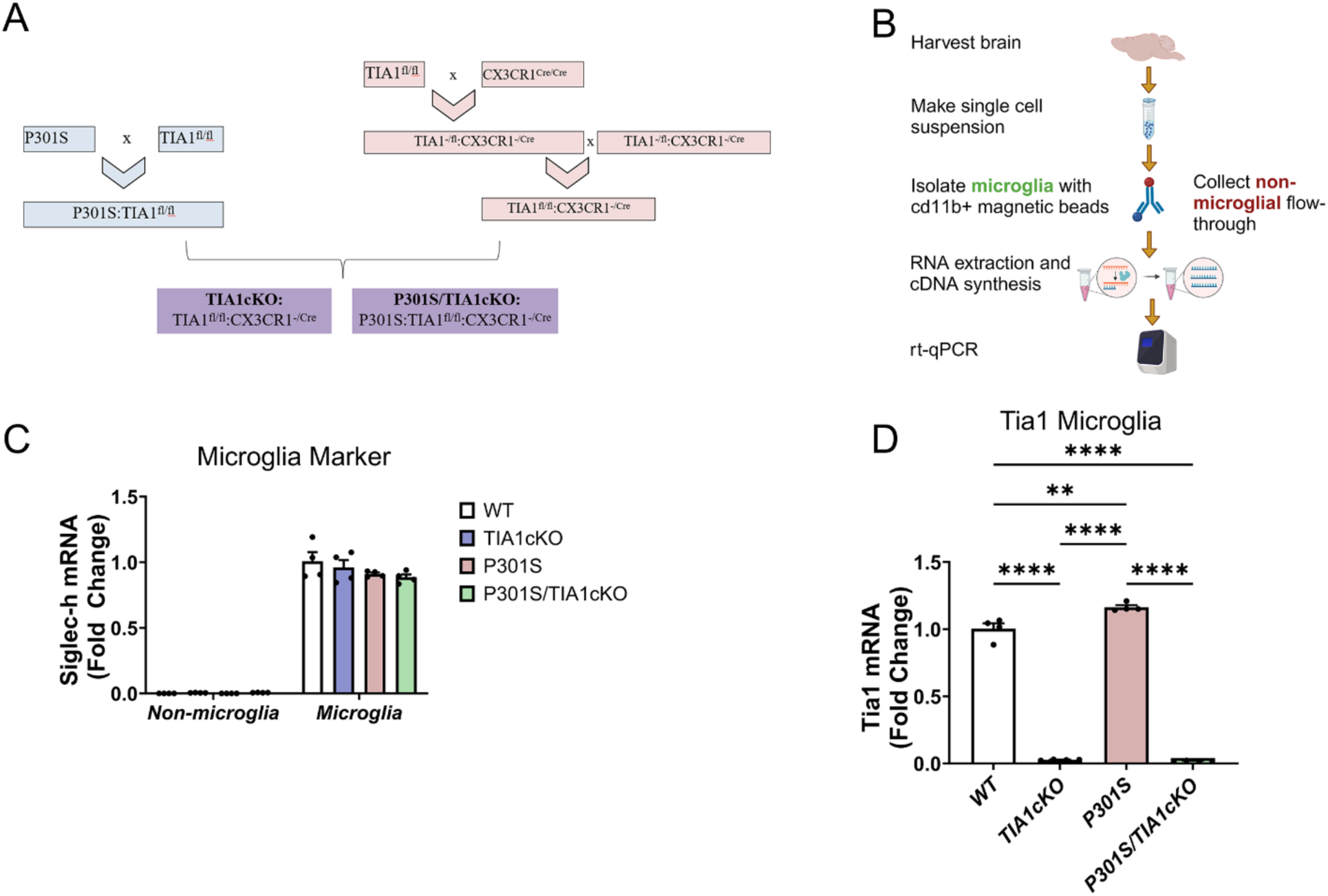
Conditional TIA1 knockout in P301S mice alters transcriptional increases of tau-specific genes. A) Breeding scheme for microglial-specific knockout of TIA1 in P301S mice. B) Schematic depicting microglia isolation using CD11b+ magnetic beads from mouse brain. C) rt-qPCR for microglial marker SIGLEC-H in CD11b+ fraction and non-microglial flow-through. D) rt-qPCR for TIA1 in microglia, N=4/genotype. One-way ANOVA (F(3,12)= 797.8, P<0.0001, followed by Tukey’s multiple comparisons with *p<0.05, **p<0.01, ***p<0.001, ****p<0.0001 Created with BioRender.com

### Microglial TIA1 deletion reduces microglial inflammatory responses

Having validated the TIA1 conditional knockout, we examined how selective loss of TIA1 in microglia affected the resulting transcriptome. For this experiment we used bulk RNA-sequencing to analyze the frontal lobe of WT, TIA1cKO, P301S, and P301S/TIA1cKO mice with CX3CR1 haplo-insufficiency (9 months, N=3/genotype) (**Figure 2A**). We identified a subset of 44 genes to be significantly upregulated in P301S mice compared to WT (p<0.05, log2foldchange>1) (**Table 1, Suppl. Tables 1**).

**Figure 2.**
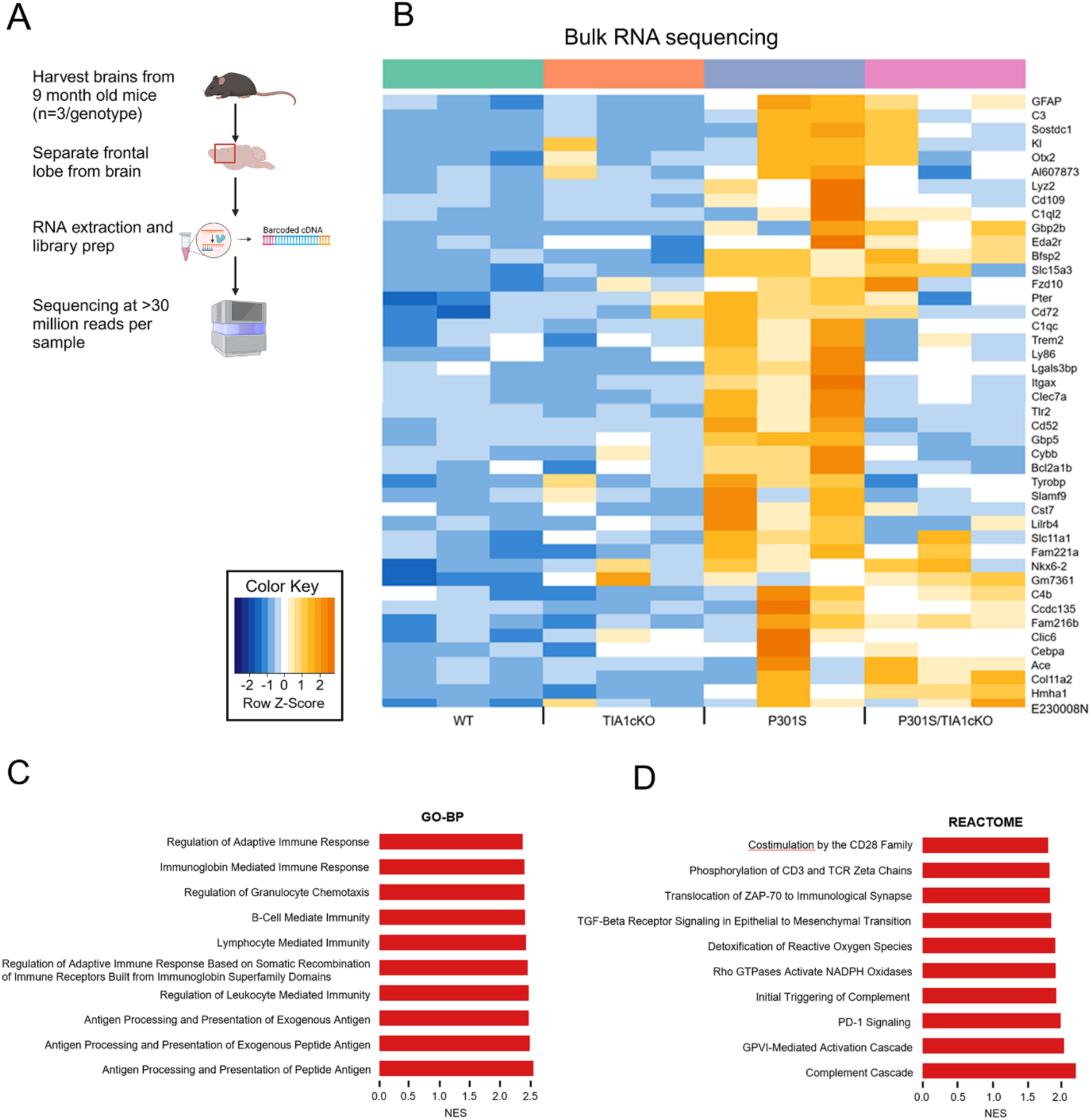
Upregulated pathways in P301S mice compared to WT. A) Bulk RNA-sequencing protocol. B) Unbiased heatmap of genes significantly upregulated in 9-month-old P301S mice compared to WT, N=3/genotype. Genes previously linked to the pathophysiology of ADRD are presented in bold orange font. C) Top 10 Gene Ontology-Biological Processes (GO-BP) pathways upregulated in P301S mice with normalized enrichment (NES) score>2. D) Top 10 Reactome pathways upregulated in P301S mice with normalized enrichment score >1.5.

**Table 1.** 44 Dysregulated genes observed by frontal lobe transcriptomics on WT, cTIA1KO, P301S, P30S/cTIA1KO mice. Upregulated: p<0.05, log2FC>1; N.C.: no change, p>0.05 or log2FC<|1|.

| Gene Symbol | cTIA1KO_vs_WT | P301S_vs_WT | P301ScTIA1KO_vs_WT |
| --- | --- | --- | --- |
| Ace | N.C. | Upregulated | Upregulated |
| Al607873 | N.C. | Upregulated | N.C. |
| Bcl2a1b | N.C. | Upregulated | N.C. |
| Bfsp2 | N.C. | Upregulated | Upregulated |
| C1qc | N.C. | Upregulated | N.C. |
| C1ql2 | N.C. | Upregulated | N.C. |
| C3 | N.C. | Upregulated | N.C. |
| C4b | N.C. | Upregulated | N.C. |
| Ccdc135 | N.C. | Upregulated | N.C. |
| Cd109 | N.C. | Upregulated | N.C. |
| Cd52 | N.C. | Upregulated | N.C. |
| Cd72 | N.C. | Upregulated | N.C. |
| Cebpa | N.C. | Upregulated | N.C. |
| Clec7a | N.C. | Upregulated | N.C. |
| Clic6 | N.C. | Upregulated | N.C. |
| Col11a2 | N.C. | Upregulated | Upregulated |
| Cst7 | N.C. | Upregulated | N.C. |
| Cybb | N.C. | Upregulated | N.C. |
| E230008N13Rik | N.C. | Upregulated | Upregulated |
| Eda2r | N.C. | Upregulated | N.C. |
| Fam216b | N.C. | Upregulated | N.C. |
| Fam221a | N.C. | Upregulated | N.C. |
| Fzd10 | N.C. | Upregulated | Upregulated |
| Gbp2b | N.C. | Upregulated | Upregulated |
| Gbp5 | N.C. | Upregulated | N.C. |
| Gfap | N.C. | Upregulated | Upregulated |
| Gm7361 | Upregulated | Upregulated | Upregulated |
| Hmha1 | N.C. | Upregulated | Upregulated |
| Itgax | N.C. | Upregulated | N.C. |
| Kl | N.C. | Upregulated | N.C. |
| Lgals3bp | N.C. | Upregulated | N.C. |
| Lilrb4 | N.C. | Upregulated | N.C. |
| Ly86 | N.C. | Upregulated | N.C. |
| Lyz2 | N.C. | Upregulated | N.C. |
| Nkx6-2 | N.C. | Upregulated | Upregulated |
| Otx2 | N.C. | Upregulated | N.C. |
| Pter | N.C. | Upregulated | N.C. |
| Slamf9 | N.C. | Upregulated | N.C. |
| Slc11a1 | N.C. | Upregulated | N.C. |
| Slc15a3 | N.C. | Upregulated | Upregulated |
| Sostdc1 | N.C. | Upregulated | N.C. |
| Tlr2 | N.C. | Upregulated | N.C. |
| Trem2 | N.C. | Upregulated | N.C. |
| Tyrobp | N.C. | Upregulated | N.C. |

These genes fall into ontology groups that include antigen presentation and the adaptive immune response in microglia, and reactome pathways that include the complement cascade and inflammatory processes (**Figure 2B**). The GO-BP pathways that were significantly upregulated in P301S mice compared to WT are involved in immune responsiveness, including the adaptive immune system, immunoglobin-mediated immunity, granulocyte chemotaxis, B-cell, lymphocyte, leukocyte, antigen processing and presentation and immune receptors (**Figure 2C**). The specific Reactome pathways that were upregulated also related to increased immune response, including CD28, CD3, programmed cell death protein 1 (PD-1) and zeta-chain-associated protein kinase 70 (ZAP-70), T and B-cell activation and activation of the complement system (**Figure 2D**).

In contrast, the P301S/TIA1cKO mice showed reduced expression for over half of the immune responsiveness genes compared to P301S MAPT mice, with many (25/44) not differing significantly from WT levels (**Figure 2B**). The top pathways increased in P301S tau mice (e.g., antigen processing and other immune functions and complement cascade) were greatly downregulated in P301S/TIA1cKO mice, showing little change compared to WT (**Figure 2C**). The GO-BP pathways show a requirement for TIA1 in full activation of the immune response normally seen in the P301S mice, and a corresponding reduction of the immune response upon TIA1 deletion. P301S/TIA1cKO mice had muted inflammatory signaling, including IL-10 production, IFNβ, IFNγ, and antigen processing and presentation (**Suppl. Figure 1A**). There was also downregulation of Reactome pathways which regulate nonsense mediated decay and translation (**Suppl. Figure 1B**). This suggests that TIA1 mediates transcription of immune related genes and deleting its activity leads to downregulation of translation.

Analysis of the TIA1-dependent microglial transcriptome revealed a striking dysregulation of tau-activated receptors and transmembrane proteins, which have been grouped under the general rubric of the “Microglial Sensome”. Out of the 100 genes known to comprise the microglial sensome, 18 genes were significantly upregulated in P301S mice but remained unchanged in P301S/TIA1cKO mice (p<0.05, vs WT, except for Itgb5) (**Figure 3A**). These included genes suspected to be involved in AD, including the Disease Associated Microglial (DAM) receptors TREM2 and CLEC7A^25,26^, and toll-like receptor 2^27^ (**Figure 3C**). The complement system is well known to be activated in AD^28–30^, which is also observed in our P301S mice (p<0.05, vs WT) (**Figure 3B**). However, many of these same complement transcripts are unchanged in PS301S/TIA1 cKO mice, including C1qa-c and complement component 5a receptor 1, C5AR1 (p<0.05, vs WT) (**Figure 3B**)^31^. The action of TIA1cKO on C5AR1 could add to neuroprotection associated with inhibition of the sensome.

**Figure 3.**
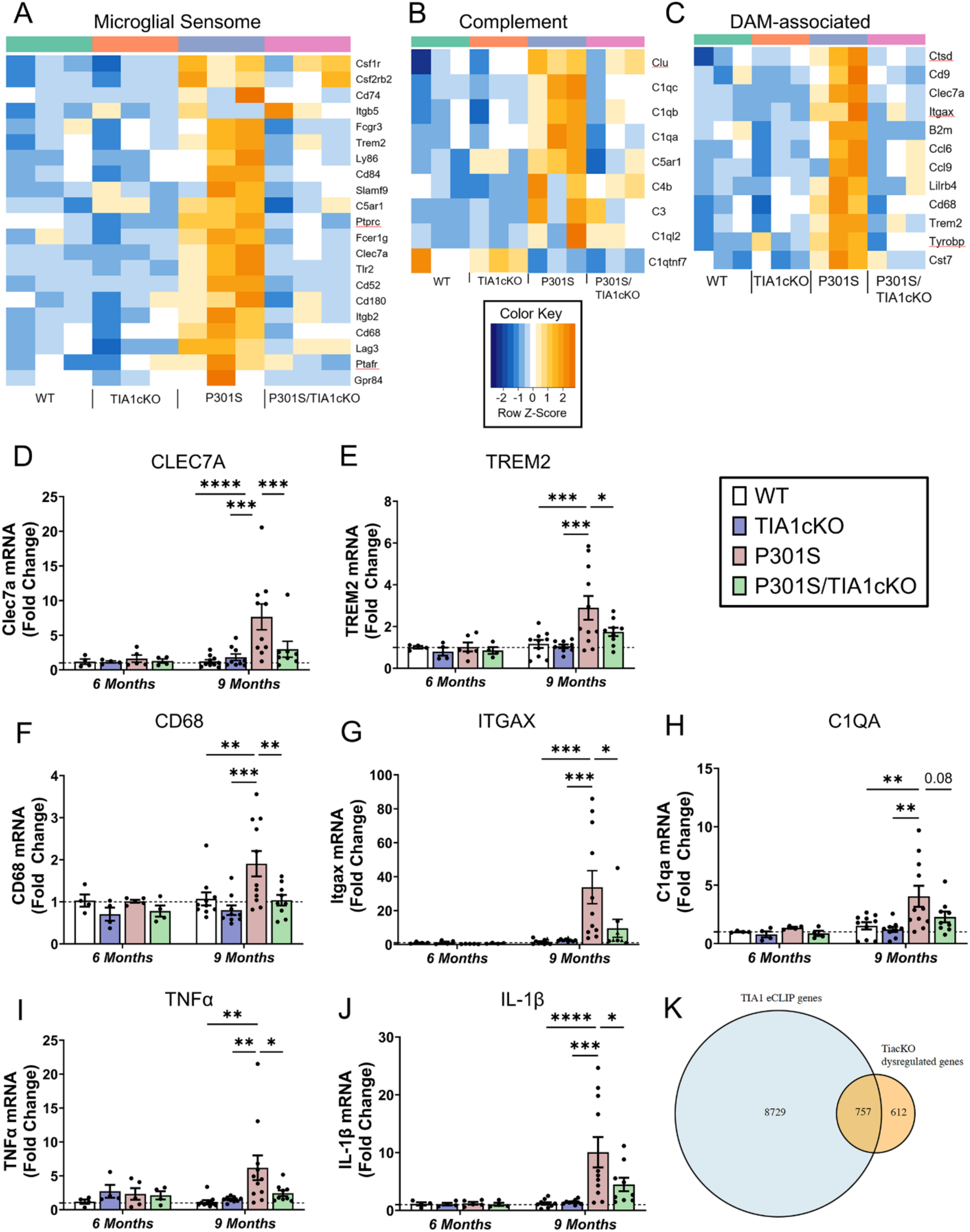
Microglial sensome response not activated in P301S/TIA1cKO mice. Heatmap depicting A) 23 microglial sensome genes, B) 12 DAM-associated genes that are significantly upregulated in P301S mice compared to WT mice, and C) 7 complement system genes, N=3/genotype. rt-qPCR amplification of D) CLEC7A (Two-way ANOVA (age F(1,46)= 5.63, p=0.022; genotype F(3,46)= 3.4, p=0.025, followed by Tukey’s Multiple comparisons), E) TREM2 (Two-way ANOVA (age F (1, 49) = 8.362, p=0.0057; genotype (F (3, 49) = 3.116, p=0.0345), followed by Tukey’s Multiple comparisons), F) CD68 (Two-way ANOVA (genotype F (3, 48) = 3.703, P=0.0178), followed by Tukey’s Multiple comparisons, G) ITGAX (9-months, P301S vs. WT Mann Whitney U=1, p <0.0001; P301S vs. cTIA1KO Mann Whitney U= 1, p<0.0001; P301S/TIA1cKO vs. P301S Mann Whitney U= 14, p<0.0121), H) C1QA (Two-way ANOVA (age F (1, 47) = 7.129, p=0.0104), followed by Tukey’s Multiple comparisons), I) TNFα (9-months, P301S vs. WT Mann Whitney U=7, p =0.0003; P301S vs. cTIA1KO Mann Whitney U= 13, p=0.0042; P301S/TIA1cKO vs. P301S Mann Whitney U= 27, p=0.095), J) IL-1β (Two-way ANOVA (interaction F (3, 48) = 2.954, p=0.04), followed by Tukey’s Multiple comparisons) in WT, TIA1cKO, P301S, and PS301S/TIA1cKO mice at 6 months (N=4-5 mice/genotype) and 9 months (N=9-11 mice/genotype). Certain values were omitted as outliers based upon 0.1% ROUT. ITGAX omitted: P301S/TIA1cKO 57.1-fold change. CLEC7A omitted: P301S 56.0-fold change and P301S/TIA1cKO 64.0-fold change. K) Overlap between TIA1 eCLIP ENCODE hits and Tia1cKO dysregulated transcripts in the 9m P301S mice. Tukey’s multiple comparisons or Mann Whitney U p-values are shown: *p<0.05, **p<0.01, ***p<0.001, ****p<0.0001

The DAM phenotype also includes some genes that are not considered part of the microglial sensome. Analysis of the DAM genes identified additional microglial transcripts that were upregulated in P301S mice but not P301S/TIA1cKO mice (p<0.05, vs WT) (**Figure 3C**). These genes include microglial endolysosomal genes, such as CD68, CST7 and CTSD.

Further RT-qPCR experiments validated the microglial-specific transcriptomic changes. Transcripts for CLEC7A, TREM2, CD68, ITGAX and C1QA were significantly upregulated in P301S mice, but not in P301S/TIA1cKO mice compared to WT (**Figure 3D-H**). Cytokines TNFα and IL-1β, important for pro-inflammatory signaling, were also increased 6.2-10.1-fold (p<0.0001 - 0.05) in P301S mice but not in P301S/TIA1cKO mice compared to WT (**Figure 3I, J**). These results validate the sequencing data and demonstrate that TIA1 cKO inhibits microglial function at multiple levels including the sensome, complement and DAM systems.

To identify RNA binding targets of TIA1, we analyzed published datasets for TIA1 enhanced crosslinking and immunoprecipitation (eCLIP) (ENCODE: ENCSR057DWB and ENCSR623VEQ). We anticipated only a partial cross-over to the brain transcriptome data because the ENCODE data was derived from clonal cell lines (HepG2 and K562 cells). Despite the difference in origins of the RNA samples, we observed over half of TIAcKO dysregulated genes (p<0.05, P301S/TIAcKO vs P301S) present in orthologous genes showing TIA1 eCLIP peak signals in both datasets, including DAM-associated genes B2m and Ctsd (**Figure 3K**). This points to direct binding of TIA1 in the activation of transcripts important for microglial response to Tau pathology and LPS.

To test whether the transcriptional dysregulation that occurred in microglia led to changes in the proportion of other abundant cell types, we deconvoluted the RNAseq data to generate cell type selective signatures using DeTREM with the GSE157827 dataset^32^. The deconvoluted data identified cell type specific signatures but showed no difference in the proportion of GABAergic neurons, astrocytes, oligodendrocytes, oligodendrocyte precursors cells, or endothelial cells between our genotypes (**Suppl. Figure 2A**). The glutamatergic neurons have selectively small variance in the P301S mice compared to other genotypes. This suggests that there is less cell type heterogeneity in these neuron populations. Dysregulation of the microglial transcriptome in TIA1cKO mice did not produce changes in the proportion of other abundant cell types.

### Microglial TIA1 is required for disease-linked morphological changes in microglia

Failure to activate the microglial sensome raised the possibility that microglial disease responses would be muted. The microglial morphologies were quantified to determine whether TIA1 cKO inhibited disease responses. Healthy microglia are large and ramified to sense different chemokines and facilitate movement. As microglia become activated, they assume an ameboid structure^15–17^. Because of this difference in morphology, the size and complexity of the microglia was measured to determine whether they are closer to a healthy, resting state or disease, activated state.

3D reconstruction of microglia visualized morphological changes caused by tau pathology (**Figure 4A**). In P301S mice, the cell size was significantly altered in response to tau pathology. Mean radius, diameter bounding circle, convex hull area, convex hull perimeter, and maximum span across the convex hull were all reduced in P301S mice compared to WT (p<0.05) (**Figure 4B-F**). Knocking out microglial TIA1 in P301S mice prevented this decrease in cell size in all metrics (p<0.05), indicating that the microglia are not activated in P301S/TIA1cKO mice.

**Figure 4.**
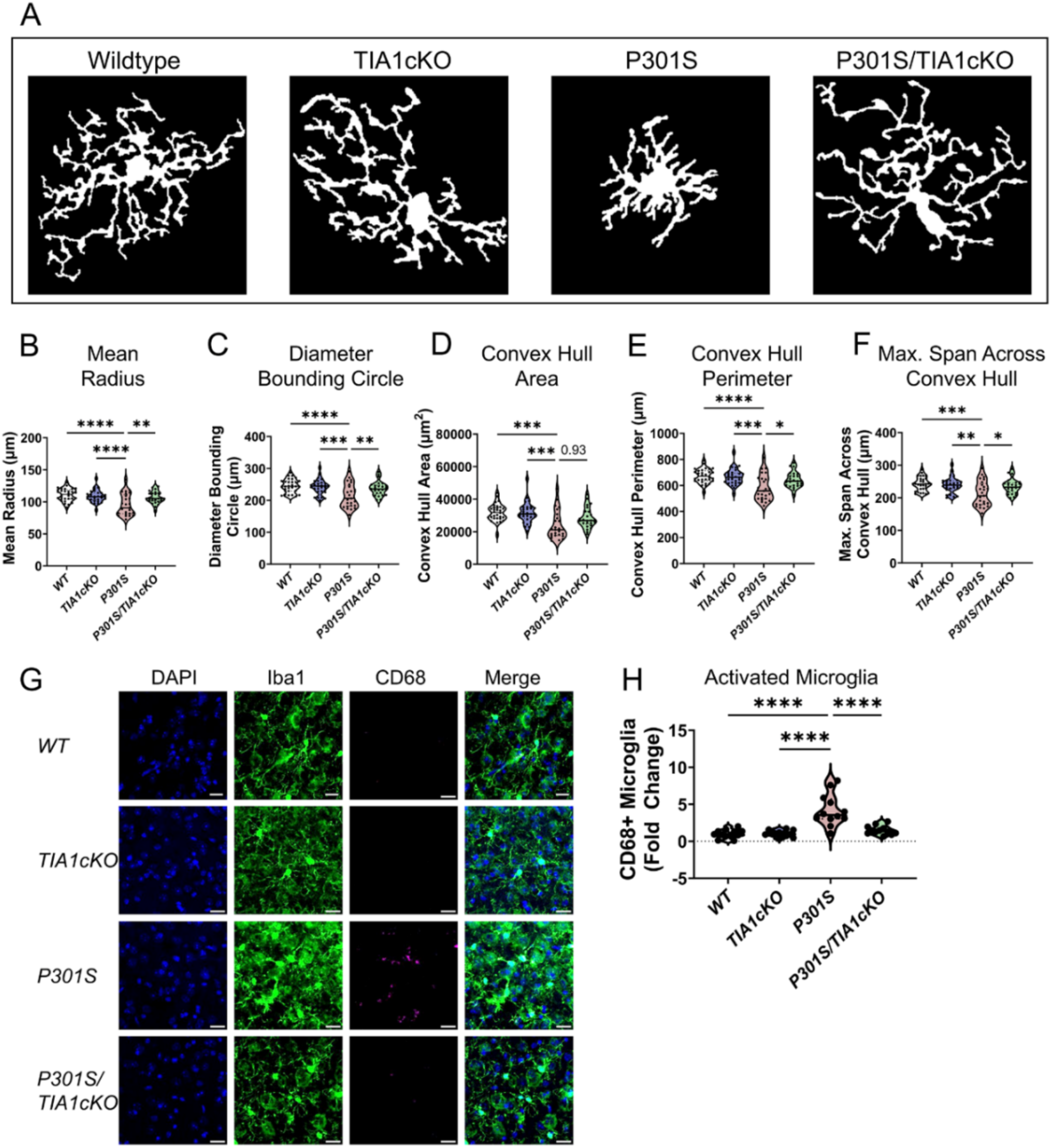
Microglia in P301S/TIA1cKO mice are not activated. A) Representative images of 3D reconstructed microglia area in 9-month old WT, TIA1cKO, P301S, P301S/TIA1cKO mice. Cell complexity measures B) mean radius (one-way ANOVA (F(3,105)=9.4, p<0.0001), C) diameter bounding circle (one-way ANOVA (F(3,105)=7.48, p<0.0001), D) convex hull area (one-way ANOVA (F(3,105)=8.08, p<0.0001), E) convex hull perimeter (one-way ANOVA (F(3,105)=4.41, p<0.0001), and F) maximum span across convex hull (one-way ANOVA (F(3,105)=6.84, p=0.0003) in WT, TIA1cKO, P301S, and P301S/TIA1cKO; N=25-30 microglia/mouse, N=5-6 mice/genotype. G) Frontal lobe was stained for DAPI, Iba1 (488), and CD68 (647) in 9-month WT, TIA1cKO, P301S, P301S/TIA1cKO mice. H) Activated microglia were quantitated as CD68+ count normalized to percent Iba1 area and fold change calculated (one-way ANOVA (F(3,44)=20.56, p<0.0001); N=2 images/mouse, N=6 mice/genotype. 10µm scale bars. All ANOVA tests were followed by Tukey’s multiple comparisons showns as: *p<0.05, **p<0.01, ***p<0.001, ****p<0.0001

Activated microglia also proliferate in response to disease. We used imaging to quantitate activated microglia in the frontal cortex using CD68 as a marker^33^ (**Figure 4G, H**); the number of CD68+ microglia were approximately 4.2-fold higher in the P301S group compared to WT (p<0.05). This increase was not observed in P301S/TIA1cKO mice (**Figure 4H**). Thus, microglial TIA1 is required for disease-linked morphological changes in microglia, consistent with the observed blockade of the microglial sensome.

### Differential responses of microglia and peripheral macrophages to LPS stimulation

Prior studies indicate that global TIA1 knockout increases peripheral inflammation in response to LPS stimulation, enabling models of autoimmune diseases, such as rheumatoid arthritis^4^. This peripheral hyper-responsiveness to inflammatory stimulation appears to contrast with the decreased responsiveness of P301S/TIA1cKO mice in the face of tau pathology. We proceeded to test whether TIA1cKO also increases responses of peripheral immune cells to LPS in our TIA1cKO mouse lines. The peripheral immune system in WT and TIA1cKO mice was stimulated with LPS, blood and brain were collected from the facial vein 3 hours after saline or 1 mg/kg LPS injection (**Figure 5A**). Cytokine levels (TNFα and IL-1β) in the periphery and the brain were measured in response to LPS injection. In the periphery we found that TIA1cKO mice had elevated TNFα and IL-1β levels compared to WT mice (**Figure 5B, C**), as has been reported previously^4^. In the brain, cytokine levels were elevated 50 to 455-fold in response to LPS but did not differ between WT and TIA1cKO injected mice (**Figure 5D, E**). Our data indicate that TIA1 inhibits the LPS response in peripheral immune cells but seemingly does not alter the LPS response in brain microglia.

**Figure 5.**
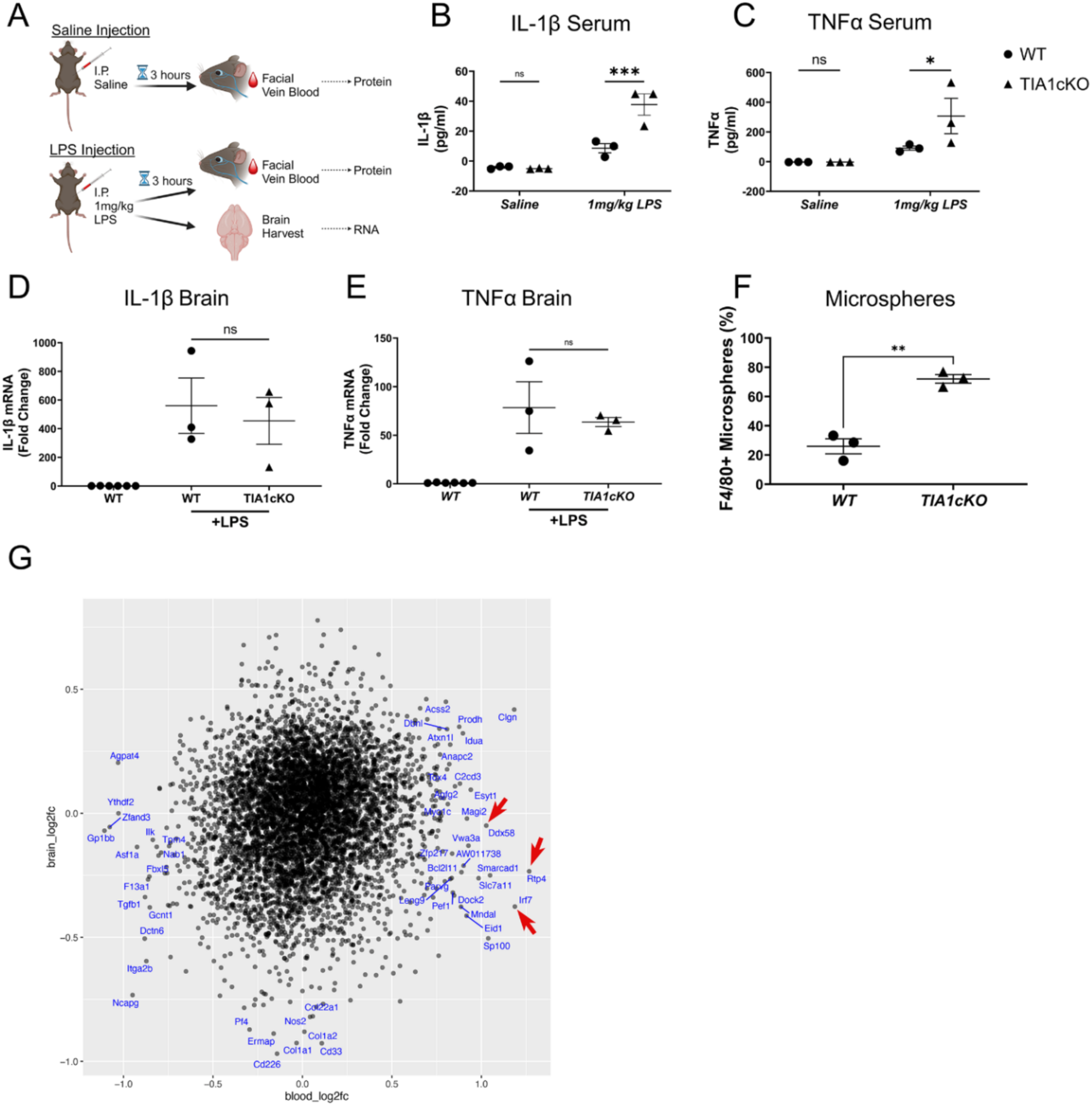
Monocyte and microglia responses to LPS stimulation and tau pathology. A) Experimental design: Blood collected from facial vein 3 hours post saline injection and post 1mg/kg LPS intraperitoneal (I.P.) injection (from the same mice), brain collected after LPS. B) IL-1β and C) TNFα serum protein amount for aged WT and TIA1cKO mice after saline or LPS injection, N=3 mice/genotype. D) IL-β and E) TNFα mRNA expression in the brain after saline or LPS injection in WT and TIA1cKO mice compared to non-LPS WT mice, N=3 mice/genotype. F) FACS data of fluorescent bead labeled peripheral macrophages (two-sample t-test (t(4)=7.78, p=0.0015)). G) Comparison of effects of TIA1 cKO on transcripts in blood and brain. The axes represent the log 2-fold change of P301S/TIA1 cKO vs. P301S tau mice for blood vs brain transcriptomes. Red arrows point to lead genes exhibiting greater differential expression in blood than brain. *p<0.05, **p<0.01, ***p<0.001, ****p<0.0001 Created with BioRender.com.

The opposing responses to LPS produced by TIA1 KO in brain microglia and peripheral macrophages prompted us to explore how TIA1 KO would affect brain penetration by peripheral macrophages. We labeled peripheral macrophages with fluorescent beads and quantified brain infiltration of these cells. TIA1cKO and WT mice were injected IV with fluorescent microbeads to label the peripheral macrophages. The mice were then stimulated with LPS (1 mg/kg, IP) and 3 hours later blood and brain macrophage levels were determined by FACS. Peripheral macrophages in the brain were elevated 5.8-fold in the TIA1cKO vs. WT mice (p=0.0015, **Figure 5F, Suppl. Figure 3C**), despite no difference in microbead labeling of monocytes (**Suppl. Figure 3A-B**). The results show an enhanced response of brain infiltrating peripheral macrophages in TIA1 cKO mice, similar to the response of peripheral macrophages. Sequencing studies were performed to explore how molecular pathways might differ between monocytes and microglia. Transcriptomes of peripheral white blood cells from TIA1cKO and WT mice were determined by bulk RNA sequencing. Interestingly, the transcriptome profiles exhibited multiple changes that could contribute to differential responses between monocytes and microglia (**Figure 5G**). Ddx58 (Rig1) and its downstream mediator, Irf7, as well as Rtp4 stood out in the monocyte transcriptome for their strong induction by TIA1 cKO in peripheral cells (p<0.05, log2foldchange>1) but not in the brain (**Figure 5G**). Ddx58 and Irf7 are known to increase activation of the interferon pathway, and enhanced Ddx58 levels increase responses to LPS^34,35^. Rtp4 is also in the interferon pathway, being induced by type IFN^36^. Thus, differential transcriptional changes in blood vs. brain contribute to the enhanced activity of peripheral macrophages compared to microglia in TIA1 cKO mice.

### TIA1 directly interacts with proteins mediating efferocytosis

TIA1 has multiple functions. As described above, as an RNA binding protein it has a strong role as post-transcriptional regulator and transcription factor. TIA1 is also well known for its role in nucleating stress granules which serve as hubs for regulating translational stress responses^7^. Besides binding RNA, each of these actions are mediated by protein-protein interactions (PPI), which could also contribute to the regulation of microglia by TIA1. We proceeded to determine the TIA1 PPIs by immunoprecipitating TIA1 from brain tissues of WT and TIA1 KO mice (global, B6.129S2(C)-Tia1^tm1Andp^/J); use of the global TIA1 KO provides a reference by allowing identification of proteins exhibiting TIA1 binding in WT but not in the TIA1 KO. We validated the anti-TIA1 antibody by demonstrating the absence of interactions in TIA1 KO (global) mice (**Suppl. Figure 4**). Next, we immunoprecipitated TIA1 from brains of WT and TIA1 KO (global) mice (N=3) followed by mass spectrometry-based identification of interaction partners (**Figure 6; Suppl Table 2**). We identified 108 interactors consistently in WT IPs (>=75%) with an >10x greater abundance relative to the TIA1KO control samples (**Fig. 6, Suppl Table 2**). GO-enrichment analysis revealed two major groups of TIA1-interacting proteins. One group of TIA1 protein interactors corresponds to the category of RNA metabolism and includes proteins regulating RNA translation and RNA binding (e.g. ribosomal proteins such as RPS11, RPS9, RPL28), consistent with TIA1’s role in stress-granule formation and RNA translation (**Suppl**. **Figure 5, Suppl. Table 3**)^7^. The second group of TIA1 protein interactors corresponds to the category of synaptic proteins, including SYP, CAMK2b, STX1A and the Alzheimer’s disease risk factor, BIN1, which has roles in both neurons and oligodendrocytes (**Suppl. Table 4**)^37–39^. These synaptic interactors are consistent with reports suggesting a role for TIA1 in synaptic function (**Suppl**. **Figure 5, Suppl. Table 4**)^40,41^.

**Figure 6.**
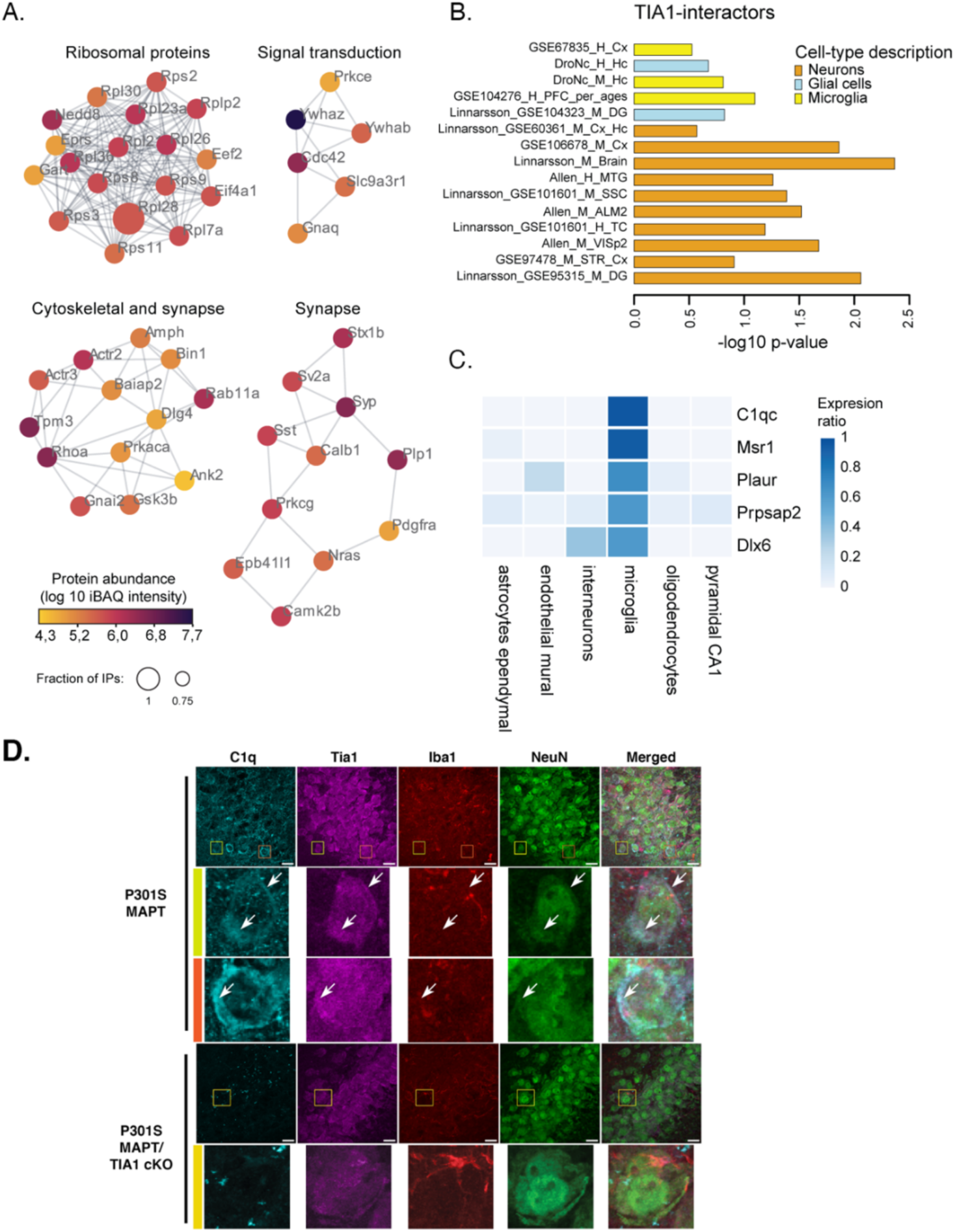
TIA1 molecular interactions. A) TIA1 IP-MS (from WT vs. Tia1^-/-^ mouse brain) revealed co-purification of 108 high-confidence TIA1-interactors. The TIA1 interactome was matched against the STRING database for known physical interactions and functional associations, and clustered into highly interconnected regions using MCODE. Only proteins detected in >=75% of wildtype IPs with an abundance >10x relative to Tia1^-/-^ were considered high-confidence interactors (WT: n= 4, Tia1^-/-^: n=3, mice were 9-months old). B) Expression weighted cell-type enrichment analysis (EWCE) using 15 datasets from publicly available human and mouse cortical tissues revealed neuronal and microglial protein signatures. C) The top 5 highest expressed proteins in microglia are part of the TIA1-interactome. Expression is based on mouse cortex scRNA-seq ^49^. D) Imaging of C1qa and Tia1 labeling in the CA3 region of the hippocampus. Colocalization of C1q and Tia1 is evident in the high magnification images in rows 2 & 3 (arrows); these images occur in NeuN positive cells indicative of neurons. Scale bar: 10 μm.

Expression weighted cell-type enrichment (EWCE) analysis revealed the presence of additional neuronal and microglial proteins in the TIA1-interactome (**Figure 6B**, **Suppl. Figure 6, Suppl. Table 5**). Notably, the top 2 microglial associated TIA1 interacting proteins were C1qc and MSR1, which are both responsible for synaptic pruning by microglia (**Figure 6C, Suppl. Table 3D**)^42–45^. Recent studies suggest that C1q interacts with RNA binding proteins ^46^. C1q was suggested to interact with ribonucleoprotein (RNP) complexes after endocytosis in neurons^46^. We proceeded to examine whether C1q and TIA1 colocalize in the brain. C1q and TIA1 were examined by immunohistochemistry in 9m P301S MAPT and P301S/TIA1cKO mice. Partial colocalization of the two proteins was observed in both microglia and neurons in the P301S mice, with neuronal C1q labeling being particularly apparent in the hippocampus, as reported previously (**Figure 6D**, C1qa shown) ^46^. Interestingly, levels of C1q were also much lower in the P301S/TIA1 cKO mice compared to the P301S mice (**Figure 6D**), suggesting that the reduced C1qa-c transcript observed in the P301S/TIA1 cKO mice (**Figure 3B**) translates to reduced C1q protein expression, including in neurons (via uptake of C1q secreted from microglia). These results suggest that TIA1 and C1q can interact in neurons, and that deletion of TIA1 in microglia downregulates C1q, including in neurons, contributing to reduced pathology and inflammation in the P301S/TIA1cKO mice.

### Microglial TIA1 is required for the development of tau pathology in P301S mice

After determining that P301S/TIA1cKO microglia do not have microglial sensome activation compared to P301S microglia, we determined whether this impacted tau pathology. In P301S mice, human mutant tau is overexpressed in neurons leading to tau aggregation. Pathological tau exhibits multiple states, including phosphorylated, misfolded, oligomeric, and fibrillar tau. We first looked at misfolded tau in the dorsal hippocampus, CA1 and CA3 regions of P301S/TIA1cKO, using the MC1 antibody (**Figure 7A-F; Suppl. Figure 7A, B**). We found that there was no difference in tau pathology at 6 months (**Figure 7B, E; Suppl. Figure 7A, B**), but strikingly by 9 months, there was a significant difference between P301S and P301S/TIA1cKO mice (**Figure 7A-F**). In the CA3 region, P301S/TIA1cKO mice have significantly less fibrillar tau and misfolded tau than P301S mice (**Figure 7B, C**). The CA1 region showed a similar significant reduction in tau pathology (p<0.05) (**Figure 7E, F**). We also examined phospho S202/205 tau using the antibody AT8 and observed similar results. For instance, immunohistochemistry for AT8 in the entorhinal cortex (EC) showed that P301S/TIA1cKO mice have significantly less phospho-tau compared to P301S mice at 9 months (**Figure 7G, H**). These results suggest that microglial TIA1 cKO reduces multiple types of tau pathology in P301S mice.

**Figure 7.**
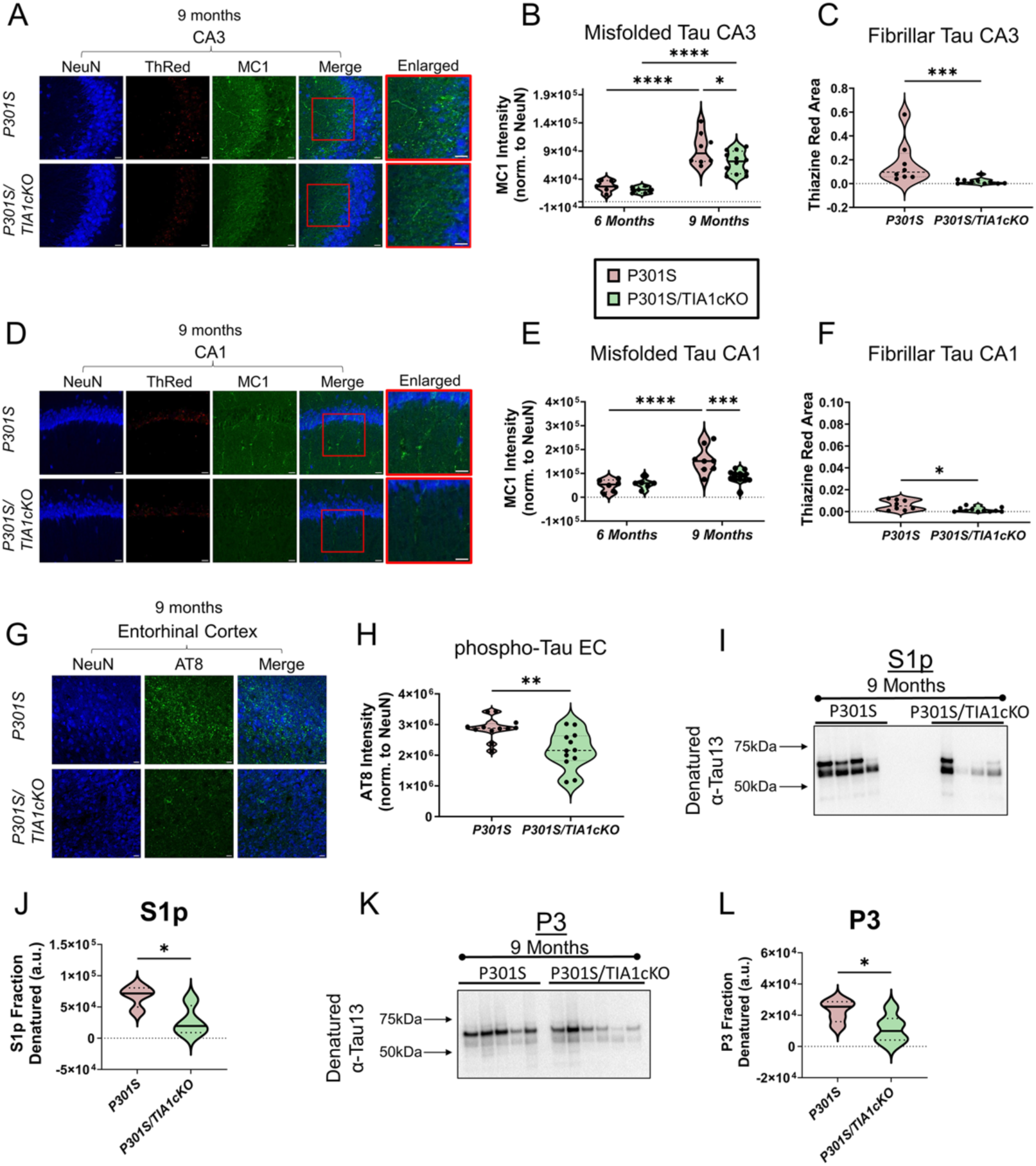
Tau pathology reduced in P301S/TIA1cKO mice. A) CA3 region D) CA1 region of 9-month-old hippocampi from P301S and P301S/TIA1cKO mice stained for NeuN (405 nm) labelling neurons, Thiazine Red (594 nm) labelling fibrillar tau, and MC1 (488 nm) labelling misfolded pathological tau. Quantification of MC1 intensity normalized to NeuN area in 6-month and 9-month-old P301S and P301S/TIA1cKO in B) CA3 (two-way ANOVA (age (F(1,28)=66.21, p<0.0001); genotype (F(1,28)=4.56, p=0.042) and E) CA1(two-way ANOVA (interaction F(1,28)=8.92, p=0.0058) regions, N=7-9 mice/genotype. Quantification of Thiazine Red are in 9-month-old P301S and P301S/TIA1cKO in C) CA3 (Mann Whitney U= 4, p=0.0005) and F) CA1 (Mann Whitney U= 15.5, p=0.016) regions. N=2 images/mouse, 6-month N=3, 9-month N=4-6 mice/genotype G) Staining of NeuN (405 nm) neurons and AT8 (488 nm) phosphorylated tau in the entorhinal cortex of 9-month-old P301S and P301S/TIA1cKO mice. N=2 images/mouse, N=6 mice/genotype H) Quantification of AT8 intensity normalized to NeuN area (two-sample t-test, t(22)=3.4, p=0.0024). I – L) Semi-nondenaturing blots with quantification of integrated lane density for S1p oligomeric tau (I, J) and P3 fibrillar tau (K, L) in 9-month-old P301S and P301S/TIA1cKO mice. N=3. 20µm scale bars; inset 10µm scale bars. *p<0.05, **p<0.01, ***p<0.001, ****p<0.0001

Next, we examined tau pathology after biochemical fractionation of brain lysates. Tau from the brains of 9-month-old P301S and P301S/TIA1cKO mice were separated into oligomeric (S1p) and fibrillar (P3) tau. These tau species were visualized on semi-nondenaturing blots to highlight the higher molecular weight tau species in each of these fractions (**Suppl. Figure 8A-D**). The S1p and P3 fractions were then run on a regular denaturing immunoblot for quantification (**Figure 7I, J**). Both pathological species exhibited significantly decreased tau in P301S/TIA1cKO mice compared to P301S mice (p<0.05) indicating that P301S/TIA1cKO mice have decreased tau pathology (**Figure 7K, L**). Thus, microglial TIA1 is important for the progression of tau aggregation in P301S mice.

## Discussion

Here, we report that selectively knocking out TIA1 in the microglia of PS19 MAPT P301S mice prevents the activation of microglia in response to tau pathology, which points to an important role for TIA1 in stimulating the neuro-inflammatory response. The reduced microglial activation was accompanied by decreased phosphorylated, misfolded, oligomeric, and fibrillar tau aggregation. The inflammation potentiating action of TIA1 in the brain contrasts with the inflammation suppressing actions of TIA1 in peripheral macrophages, which has been reported previously^4^. Thus, TIA1 appears to function within a hub that controls the phenotypes of microglia and peripheral macrophages.

The actions of TIA1 in microglia and peripheral macrophages occur on at least two levels, transcript-protein and protein-protein interactions. Regulation of transcript levels by TIA1 was evident from the sequencing and qPCR data. which corroborates previous research on TIA1-like protein’s role in transcription regulation^47,48^. Transcripts associated with the microglial sensome, complement system and endolysosomal system were strongly increased in brains of P301S MAPT mice compared to WT mice. In contrast, such disease-linked increases were absent in P301S/TIA1cKO mice compared to WT mice. Of these regulated transcripts, several were previously shown to directly interact with TIA1. TIA1 PPI that might contribute to regulation of microglia are delineated by the IP-MS data, which reveal a robust interactome for TIA1 in the brain. A prominent component of the TIA1 brain PPI is C1q, which is a microglial protein expressed and secreted upon microglial activation. C1q is produced by microglia, secreted and taken up by neurons^46^. The C1q complex consists of 6 heterotrimeric subunits, each containing one C1qa, C1qb and C1qc chain^49^. C1qc is the most consistent and abundant TIA1 protein interactor; C1qa protein was also detected in the TIA1 protein interactome with >10x endogenous abundance. Levels of C1qa protein were reduced in the P301S TIAcKO hippocampus, which is consistent with the reduced C1qa-c transcript expression in P301S/TIA1 cKO mice (compared to P301S MAPT), reflecting transcriptional regulation by TIA1. The interaction of TIA1 with C1q is supported by recent paper showing age-related increases in C1q interactions with ribonuclear protein complexes, which include RNA binding proteins^46^. C1q is also one of the most prominent proteins upregulated in the AD brain and in mouse models of AD^49,50^, where it is thought to mediate synaptic pruning by microglia^22,42,43,51^. Conversely, microglial-selective TIA1 deletion might inhibit the C1q pathway, reduce microglial activation, synaptic pruning and dampening the process of neurodegeneration.

The mechanism through which TIA1 might differentially regulate microglia and peripheral macrophages is suggested by the pattern of transcript expression shown in our transcriptome studies. This finding complements recent studies suggesting differential biology of peripheral macrophages and microglia^52^. Prior literature and our own studies indicate that LPS stimulation of TIA1 knockout mice produces a robust peripheral inflammatory response^4^. In the periphery, LPS binds directly to TLR4 on macrophages. TLR4 activates the NF-κβ and IRF3 transcription factors producing a cytokine storm that releases interleukins and other cytokines into the circulatory system, producing systemic inflammation^53–56^.

Importantly, some key transcriptional changes occurring in peripheral macrophages did not occur in brain microglia. LPS stimulation selectively increased Ddx58 (Rig1) and Irf7 in peripheral macrophages, which act in the interferon pathway; Rtp4, which is induced by the interferon pathway, was also selectively increased in peripheral macrophages^34,35,57^. Deletion of TIA1 in macrophages also strongly increased Chil3 (produced by macrophages and neutrophils) and moderately increased TLR4, Trem2 and TyroBP in monocytes but not in microglia; these all would further potentiate the inflammatory response in peripheral macrophages^58–61^. Thus, differential regulation of key transcripts (Ddx58, Irf7, Rtp4, Chi3, Tlr4, Trem2 and TyroBP) contributes to the divergent regulation of microglia and monocytes by TIA1 deletion. Differential regulation of TIA1 mediated protein signaling pathways also might contribute to the divergent actions of microglia and macrophages. Our results show that TIA1 acts in the C1q pathway, which regulates synaptic pruning in the brain but clotting in the periphery^22,42,43,51^. The disparate functions of C1q in brain and periphery might thus contribute to the differential actions of TIA1 in microglia and peripheral macrophages. The results presented above demonstrate the multi-modal actions of the RNA binding protein TIA1, highlighting roles in regulation of the microglial sensome, complement and possibly synaptic biology. This study also suggests that TIA1 functions as part of a hub that controls the phenotypes of microglial and peripheral macrophages allowing the two cell types to generate opposing responses to inflammatory stimuli. These results show that the biology of TIA1 extends well beyond its role in nucleating stress granules, exerting broader roles in the brain, regulating microglial and neuronal functions.

## Acknowledgements

Funding for BW: (AG059925, AG050471, AG050471, R01AG080810, AG056318, AG064932 (shared with AE), AG061706 (shared with AE), and UO1AG072577), the BrightFocus Foundation and the Alzheimer Association. Funding for HL: H.L. was supported by grants from the Mayo Clinic Center for Biomedical Discovery, the Mayo Clinic Center for Cell Signaling in Gastroenterology (NIH: P30DK084567), the Glenn Foundation for Medical Research, the Eric & Wendy Schmidt Fund for AI Research & Innovation and the National Institutes of Health (NIH; U19AG74879, P50CA136393, R03OD038392). Funding for JS: JMS was supported by national funds through the Portuguese National Funding Agency (FCT) 2021.00204.CEECCIND, UIDB/50026/2020 and UIDP/50026/2020; ICVS Scientific Microscopy Platform, member of the national infrastructure PPBI - Portuguese Platform of Bioimaging (PPBI-POCI-01-0145-FEDER-022122). Funding for AOR: Rainwater Foundation

## Author contributions

C.J.W. conceived, performed and analyzed experiments and wrote the original draft; S.J.F., performed and analyzed experiments; ALC generated the mice and performed experiments; SP performed experiments; CZ analyzed experiments; JTMA performed and analyzed experiments; GZP performed and analyzed experiments; RR performed experiments; KKJ performed experiments; THT performed experiments performed experiments; AT performed experiments, ZW performed and analyzed experiments, JP performed experiments, IS provided expertise and edited the manuscript; EM designed and supervised experiments, provided funding and edited the manuscript; JS, designed and supervised the experiments and edited the manuscript; B.W. conceived, analyzed, wrote the original draft, edited, supervised and provided funding for the project.

## Materials and Methods

### Animals

All use of animals was approved by the Boston University Institutional Animal Care and Use Committee. Mice were fed *ad libitum* and housed in a facility with 12 hour on/off light cycles. All mice used in this study are on a C57BL/6 genetic background. P301S mice overexpressing human P301S tau with a prion promoter (B6;C3-Tg(Prnp-MAPT*P301S)P301SVle/J, strain #: 008169) and C57BL/6J mice (strain# 000664) were purchased from Jackson Laboratories. TIA1^fl/fl^ mice (C57BL/6NTac-Tia1tm1a(KOMP)WTsi/WTsi, EM: 11228) were purchased from a European Conditional Mouse Mutagenesis Program. CX3CR1 *cre* mice (B6J.B6N(Cg)-CX3CR1tm1.1(cre)Jung/J, strain #: 025524) were purchased from The Jackson Laboratory. To achieve our genotype of interest, P301S:TIA1^fl/fl^ mice were bred to TIA1^fl/fl^:CX3CR1^Cre/Cre^ mice to produce P301S:TIA1^fl/fl^:CX3CR1^-/Cre^ mice (referred to as P301S/TIA1cKO).

Mice were sacrificed with an overdose of ketamine/xylazine cocktail and perfused with Dulbecco’s phosphate-buffered saline (DPBS) (w/o Ca^2+^ or Mg^2+^). The brain was harvested and hemispheres separated. One hemisphere was dissected to remove the frontal lobe and then the two pieces flash frozen and stored at −80 °C. The other hemisphere was drop-fixed for 1.5-2 hours in 4% paraformaldehyde then stored at 4 °C in PBS with sodium azide. For CD11b+ microglia isolation, the brain was processed fresh.

Global *Tia1*^-/-^ mice (B6.129S2(C)-Tia1^tm1Andp^/J) were generated by Anderson and colleagues and obtained from Harvard University, Dana Farber Cancer Institute^4^; these mice had previously been backcrossed for 10+ generations to the C57BL/6 J inbred strain. These mice were raised at the University of Minho (Portugal); use of the animals was approved by the U. Minho Institutional Animal Care and use Committee.

### Microglia isolation

Single cell suspensions were made from fresh whole brain of mice perfused with PBS. Two mouse brains for each genotype were combined for microglia isolation. Tissue was dissociated using the Neural Tissue Dissociation Kit (P) (Miltenyi Biotec, cat# 130-092-628) per the protocol for Manual dissociation for gene expression profiling. The single cell suspensions were then treated with Debris Removal Solution (Miltenyi Biotec, cat# 130-109-398) to remove myelin debris. CD11b Microbeads (Miltenyi Biotec, cat# 130-093-636) were used to bind CD11b^+^ microglia. Isolated microglia and flow-through were collected and their RNA immediately extracted using the Qiagen RNeasy Plus Mini Kit (cat# 74134) RNA was eluted in 30 µl of RNase-free water and stored at −80 °C.

### RNA extraction

Frontal lobe (∼25 mg) of fresh frozen mouse brains were moved into 1 mL of TRIzol and lysed using pre-filled sterile bead mill tubes (Fisher, cat# 15-340-153) with Tissue Homogenizer. The lysate was allowed to incubate on ice for 5 minutes, then 200 μL of chloroform was added. The samples were vortexed on medium setting for 15 seconds then allowed to incubate on ice for 3 minutes. The samples were centrifuged at 12,000 g for 15 minutes at 4 °C to phase separate the RNA. The aqueous RNA layer from removed and processed with the Qiagen RNeasy Plus Mini Kit (cat# 74134) per the protocol. RNA was eluted in 30 µl of RNase-free water and stored at −80 °C.

### Reverse transcription quantitative polymerase chain reaction

RNA was converted to complementary DNA (cDNA) using the High-Capacity cDNA Reverse Transcription Kit (Applied Biosystems, cat# 4368814) with the addition of RNase Inhibitor (Promega, cat# N2615) per the protocol. Taqman primer probes were used to amplify transcripts. The Taqman Assays used were: SIGLECH-H (Mm00618627_m1), TIA1 (Mm01183616_m1), C1qa (Mm00432142_m1), IL-1β (Mm00434228_m1), TNFα (Mm00443258_m1), TREM2 (Mm04209424_g1), CLEC7A (Mm01183349_m1), ITGAX (Mm00498701_m1), CD68 (Mm03047343_m1), glyceraldehyde 3-phosphate dehydrogenase (GAPDH) (Mm03302249_g1). Samples were run in triplicate or quadruplicate. Quantitative PCR was performed on a Quantstudio 12K Flex qPCR system and cycle threshold (Ct) values were compared by fold change of ΔΔCt. The raw Ct values were first normalized to GAPDH (ΔCt), then normalized to an average of wildtype ΔCt (ΔΔCt), finally the values were converted to fold change.

### Image acquisition for microglial morphological analysis

For the analysis of microglial morphology, blinded researchers imaged two coronal brain sections per animal (n = 5-6 per genotype) for the region of interest (CA1 subregion of the hippocampus). The Olympus Confocal FV3000 (Tokyo, Japan) laser scanning microscope with a resolution of 1024 × 1024 px using a 40× objective (UPlanSApo, N.A. 0.95; dry; field size 318.20 × 318.20 μm; 0.31 μm/px) was used to obtain all Z-stacked images. The acquisition settings were the following: scanning speed = 2 μm/px; pinhole aperture = 120 μm; ionized calcium-binding adapter molecule 1 (Iba-1), excitation = 590 nm, emission = 618 nm; in a 3-dimensional scenario (X, Y, and Z axes).

### Image analysis and microglial morphological data acquisition

Imaging analysis of microglia was performed blinded. The ImageJ software (v1.54f; National Institute of Health, Bethesda, MD, USA) was used on Z-stacked from sections of the brain region of interest (CA1). For morphological analysis, a semi-automatic method adapted from^62^ was used, that was developed by Campos et al^63^.

Binary images were created to acquire fractal and skeleton data. Following the stacking of 3D volume images with maximum intensity projection and conversion into 8-bit, the two-channel image was split to isolate the Iba-1 channel, and converted to grayscale to best visualize all positive staining where there were adjusted brightness and contrast. Subsequently, the unsharp mask filter was used to further increase contrast and the despeckle filter was applied to eliminate noise, with the threshold option being fine-tuned as required before finally removing any outliers with the remove outliers filter (radius 2.0 px, threshold 50). Using the rectangle tool of 250 x 250 px, five cells per image were chosen. Post-selection, the morphology of the cells was refined using the paintbrush tool, resulting in binary single microglial-cell images.

The data extraction, pre-processing and organization of morphological features of single microglial cells associated with cell complexity (FracLac plugin) and ramification [Analyze Skeleton (2D/3D) plugin] was done with the MorphData plugin^63^. The total number of microglial cells used was 109 (30 from WT mice, 29 from TIAcKO, 25 from P301S, and 25 from P301S/TIA1cKO mice).

### Bulk RNA sequencing

RNA was isolated from the frontal lobe of n=3 mice/genotype using protocol listed above. For the LPS experiment, 3 hours after LPS administration facial vein blood was collected and the brain harvested, n =4/genotype. RNA was extracted from clotted blood and frontal lobe tissue per protocol above. The sequencing libraries were prepared using SMARTer stranded Total RNA-Seq Kit v2-Pico Input Mammalian Kit following manufacturer instructions (Takara Bio USA, Inc). Briefly, 10ng of total RNA was subjected to fragmentation followed by first-strand cDNA synthesis. Subsequently, the libraries were barcoded with dual indexes and purified using AMPure beads. Depletion of ribosomal RNA was performed followed by another round of purification and library amplification. The cDNA libraries underwent validation through the Agilent Bioanalyzer and were then subjected to sequencing on an Illumina NextSeq 2000, yielding a read depth of >30 million reads per sample.

Fastq files of paired-end reads were mapped to the UCSC mm10 reference genome using STAR version 2.6.0a in 2-pass mode^64^. Gene-level counts were obtained with featureCounts version 1.4.6^65^. Differential expression analysis was done using DESeq2 version 1.28.1^66^. Gene Set Enrichment Analysis (GSEA) was performed as using the GSEA Java package^67^ against the mouse gene sets from Enrichment Map^68^. Over-representation analysis was done using Fisher’s exact test. Significance was determined using the unadjusted p-value at an alpha of 0.05. To run DeTREM deconvolution, a reference single-cell RNA-seq dataset GSE157827^69^ was downloaded from the Gene Expression Omnibus and was mapped to mouse homologs according to the Mouse Genome Database^70^. DeTREM deconvolution was then done using R package DeTREM^32^ on gene-level counts, with the reference dataset in mouse homologs. Deconvolution using seq-ImmuCC^71^ gene signatures was done with CIBERSORTx^72^.

### Brain homogenization and tau fractionation

Fresh frozen brains were weighed and homogenized with 4X w/v HSAIO buffer (50 mM Tris base, 274 mM NaCl, 5 mM KCl, pH 8.0) with protease (Pierce, cat# A32955) and phosphatase (Roche, cat# 04906845001) inhibitors using a motorized mortar and pestle. Equal volumes of brain lysate were used for tau fractionation. The remaining brain lysate was lysed further by addition of 0.2% NP40 detergent and chelating agent 2 mM EDTA and then spun at 15,000 g at 4 °C for 15 minutes and the supernatant aliquoted and stored at −80 °C.

For tau fractionation into soluble oligomeric tau and insoluble fibrillar tau, an equal volume of brain lysate was spun down at 28,000 rpm at 4 °C for 20 minutes in an ultracentrifuge with rotor TLA-55k. The soluble supernatant was removed and pelleted at 55,000 rpm for 45 minutes at 4 °C to isolate oligomeric tau while the insoluble pellet was used for fractionation of fibrillar tau. This insoluble pellet was homogenized in a high salt-sucrose Buffer B (10 mM Tris, pH 7.4, 0.8 M NaCl, 10% Sucrose, 1mM EGTA) and spun down at 22,000 rpm at 4 °C for 20 minutes. An equal volume of supernatant was removed from each sample and incubated with 1% Sarkosyl for 1 hour at 37 °C shaking at 300 rpm. The Sarkosyl insoluble tau (fibrillar tau) was then pelleted by spinning at 55,000 rpm for 1 hour at 4 °C. The tau fractions were resuspended in TE Buffer (pH 8.0), aliquoted, and stored at −80 °C.

### Immunoblots (denaturing and semi-nondenaturing)

For denaturing western blot, brain lysate protein was quantified with BCA (Pierce, cat# 23224 and 23223) per the protocol. Samples were combined with 4X LDS Sample Buffer (Invitrogen, cat# B0007), 10X reducing agent (Invitrogen, cat# B0004), and brought to equal volume with water. Samples were denatured for 5 minutes at 95 °C and then loaded onto a Bolt 4-12% Tris-Bis Gel with MOPS running buffer (Invitrogen, cat# B0001).

For semi-nondenaturing western blot, tau fractions (oligomeric or fibrillar) were combined with 4X LDS Sample buffer and brought to equal volume with water. The samples were then loaded on Blot 4-12% Tris-Bis Gels with MOPS running buffer.

All gels were transferred to PVDF membranes (Invitrogen, cat# IB24002) using the iBlot 2 Dry Blotting system. Semi-nondenaturing gels were transferred at 20V for 2 minutes, 23 V for 4 minutes, 25 V for 3 minutes. Denaturing gels were transferred using P0 template (20V 1 minute, 23 V 4 minutes, 25 V for 3 minutes). Membranes were blocked in 5% non-fat milk for 1 hour at room temperature than incubated overnight with primary antibodies. Primary antibody used was Tau13 (Dr. Peter Davies, 1:10,000 - 1:50,000).

### Immunohistochemistry

Brains were transferred to 30% sucrose and stored at 4 °C for at least 72 hours and up to 3 weeks. Brains were sectioned at 35 µm slices on the cryostat and stored in DPBS with 0.01% sodium azide at 4°C. For long term storage, brain sections were stored in cryoprotectant at −20 °C. Two brain sections per mouse were selected from bregma 1.5 to 2.0 mm (frontal lobe), −1.5 to −2.0 mm (dorsal hippocampus), and −2.8 to −3.3 mm (ventral hippocampus) for IHC.

Free floating sections were permeabilized with 0.25% Triton X-100 TBS for 15 minutes rocking, then blocked for 2 hours rocking, at room temperature in 5% Bovine Serum Albumin, 5% Donkey Serum in 0.1% Triton X-100 TBS (TBST-0.1%). Sections were then incubated overnight in primary antibody in TBST-0.1%, rocking, at 4 °C, with photobleaching. Primary antibodies used were: NeuN (EMD Millipore, cat# ABN91, 1:1000), MC1 (Dr. Peter Davies, 1:300), AT8 (ThermoFisher, cat# MN1020, 1:1000), Iba1 (Wako, cat# 019-19741, 1:1000), CD68 (Bio-Rad, cat# MCA1957, 1:600), C1q (Abcam, cat# 7H8, ab11861, 1: 200), TIA1 (Abcam, cat# ab140595, 1: 200). The following day, sections were washed twice with TBST-0.1% for 10 minutes, rocking, at room temperature, then incubated with appropriate Alexa Fluor IgG secondary antibodies (1:800) for 1 hour, rocking, at room temperature. Sections were washed once and then, if appropriate, DAPI (1:10,000) or Thiazine Red (1:1000) were added for 15 minutes rocking at room temperature. Sections were then washed twice more and mounted onto Superfrost coverslips (Fisher Scientific, cat# 12-550-15). After sections dried, coverslips were added using ProLong Gold Antifade (Cell Signaling Technology, cat# 9071S). Slides were left to dry at room temperature, then they were sealed with clear nail polish and stored at −20 °C. Z-stack images were acquired by Keyence BZ-X700 fluorescent microscope or Zeiss LSM700 confocal microscope for quantification and representative images by Zeiss LSM700 confocal microscope.

### Image analysis

Fiji (ImageJ) was used for all image analysis. For immunohistochemistry, Z-stacks were projected as maximum or average pixel intensity. The integrated density was quantified for tau antibodies, MC1 and AT8, and normalized to the NeuN area determined through thresholding of the NeuN positive neurons. Thiazine Red positive aggregates were counted with Analyze Particles by first adjusting the threshold (37-255). Iba1 and CD68 staining was quantified by normalizing the CD68 particle count to the Iba1 Area. CD68 was counted with Analyze Particles by first subtracting background (50 pixels) then thresholding (28-255). Fold change to WT was calculated. For denaturing western blots, integrated densities were calculated for each band.

### Lipopolysaccharide and ELISAs

Blood was collected from the facial vein from mice aged 9 – 11 months at two timepoints: post-saline injection and post-LPS injection. These blood collections were separated by at least 5 days to allow the veins to heal. Blood was collected 3 hours post either intraperitoneal saline or LPS (1 mg/kg) injection. The blood was collected in 1.5 mL Eppendorf tube and stored at 4 °C for 16-24 hours to allow the blood to clot. The serum was then separated by centrifugation at 3,000 g for 15 minutes at 4 °C and saved at −80 °C. BCA was performed to quantify serum protein concentration. 150 μg of serum protein was analyzed using ELISA for IL-1β (R&D Systems, cat# MLB00C) and 350 μg of serum protein was analyzed on ELISA for TNFα (R&D Systems, cat# MTA00B) per the protocols. Samples were run in duplicate. The mice were sacrificed 3 - 6 hours post LPS injection, perfused with DPBS, and their brains flash frozen and stored at −80 °C. RNA was then extracted from the frontal lobe and used to quantify cytokine levels through rt-qPCR.

### Fluorescence activated cell sorting (FACS)

Mice (N=3/genotype) were injected twice via tail vein with 125μL Fluoresbrite® Polychromatic Red Microspheres 0.5µm (cat# 19507-5) diluted 1:25 with PBS 24-hours apart. Twenty-four hours after the second tail vein injection, mice were injected with 1m/kg LPS intraperitoneally. Blood and brain were harvested 3 hours post-LPS. Cardiac blood collected into a tube with 1μL 10% k-EDTA per 100μL blood. Ammonium chloride solution (Stemcell Technologies, cat# 07800) was added to the blood at a 9:1 ratio. The cell suspension was mixed thoroughly and incubated on ice for 10 minutes. The cells were then washed twice in FACS buffer (DPBS with 1% fetal bovine serum) then resuspended at 1E7 cells/mL. The brain was processed using the Neural Tissue Dissociation Kit (P) (Miltenyi Biotec, cat# 130-092-628) manual dissociation protocol. In brief, tissue was dissociated mechanically and enzymatically before being made into a single cell suspension using a 70μm cell strainer. Tissue debris was removed using the Debris Removal solution (Miltenyi Biotec, cat# 130-109-398) per the protocol. The cells were counted on a Countess Automated Cell Counter and resuspended at 1E7 cells/mL in FACS buffer. The cells were then incubated with 1:50 F4/80:Alexa Fluor® 488 (Bio-rad, cat# MCA497A488T) for 30 minutes at 4°C, rocking. Cells were then washed with FACS buffer and stained with LIVE/DEAD™ Fixable Violet Dead Cell Stain (ThermoFisher Scientific, cat# L34963) per the protocol. Cells were stored overnight at 4°C in eBioscience™ IC Fixation Buffer (ThermoFisher Scientific, cat# 00-8222-49). Flow cytometric analyses was performed on a Beckman Coulter MoFlo Astrios and analyzed on FlowJo v10.10.0 software.

### Immunopurification

Wildtype and *Tia1*^-/-^ mice (B6.129S2(C)-Tia1^tm1Andp^/J) hippocampi were homogenized in protein extraction buffer (10 mM Tris-HCl pH7.5 (Bioworld, 21420063-2), 150 mM NaCl (Sigma-Aldrich, S6546), 0.5 mM EDTA (Thermo Scientific, AM9260G), 0.5% N-P40 (EMD Millipore, 492016), 1% CHAPS (Fisher Chemical, BP5715), supplemented with Phosstop phosphatase inhibitor (Roche, 4906845001) and Halt protease inhibitor cocktail (Thermo Scientific, 78430)) using a dounce homogenizer, 12 strokes. After incubation (1h, 4°C, rotating), samples were spun down (12000 xg, 10 min, 4°C) and protein concentrations in were determined using BCA. Lysates were diluted to 1μg/μL with extraction buffer, and 200 μL was incubated with 2 μg antibody (ab140595) overnight at 4°C, rotating. The following day, samples were incubated for 1h at 4°C, rotating, with 20 μL Dynabeads (Invitrogen, 10008D) prewashed in washing buffer (10 mM Tris-HCl, 150 mM NaCl, 0.5 mM EDTA, 0.1% Triton-X). Samples were pulled-down, washed 1x with washing buffer and twice with 50 mM Ammonium bicarbonate (ABC) before on-bead digestion with 1 μg trypsin dissolved in ABC (60 μL, 0.16 μg/μL). After incubation overnight at 37°C, rotating, samples were desalted using Pierce C18 spin columns (Thermofisher, 8987), dried in Speedvac and stored at −20°C until mass spec analysis.

### UHPLC-MS/MS and Data analysis

Peptides were re-dissolved in 15 μL of 0.1% formic acid in water, and loaded onto a reverse-phase trap (75 μm i.d. × 2 cm, Acclaim 3 μm 100 Å PepMap100-C18 resin, ThermoScientific, Waltham, MA, USA). Next, peptides were separated on an EASY-Spray™ PepMap™ Neo UHPLC Column column (75 μm i.d. × 50 cm, 2 μm, 100 Å; **<u>ES75500PN</u>**, ThermoScientific) with Ultrahigh Performance Liquid Chromatography (UHPLC) on a Neo Vanquish system (ThermoScientific), using a linear gradient of increasing acetonitrile concentration from 0 to 3.2% in 6.5 min, and to 31.2% in 103.5 min with a flow rate of 0.3 μL/min. Peptides were electrosprayed into an Orbitrap Exploris 480 mass spectrometer (ThermoScientific), operated in a data dependent manner with one MS (m/z range from 150 to 1500) followed by MS/MS with a normalized AGC target of 300% in the MS1 domain and 50% in the MS2 domain, an intensity threshold of 5.0e4 and an exclusion window of 45 sec. The resulting MS/MS spectra were searched against the Mouse proteome (UP000000589_10090, 2024_07_17) with MaxQuant software (version 2.6.2.0). The search parameters were set to unique and razor peptides for protein quantification and digestion with trypsin. True interactors were determined based on count (# samples identified) and abundance. Proteins identified in >= 75% of wildtype IPs with >10x higher iBAQ intensity versus Tia1KO were considered true interactors. Enriched proteins were used to build a protein network in Cytoscape Version 3.10.2 and sub-clustered with the MCODE plugin. GO-enrichment analysis was performed with ShinyGO Version 0.80^73^. Expression weighted cell-type enrichment (EWCE) analysis was performed as implemented in FUMA^74^. Pre-processed single-cell RNA sequencing data from human and mouse cortical tissues were used for analysis. The cell-type enrichment matrix shown is based on Watanabe et al ^74^.

### Statistics

All statistical analyses were performed in GraphPad Prism (v10.0.3). Statistical significance was set at an α = 0.05. All data are presented as mean ± SEM. Outliers were identified using ROUT method set to Q = 0.1% and excluded from analysis, all omitted outliers are noted in the text. Column data was compared using unpaired two-sided t-test or one-way ANOVA, a significant difference prompted pairwise comparisons controlled for by Tukey post-hoc test. Grouped data was compared using two-way ANOVA with comparisons within each age group, Tukey’s post-hoc test was used for significant differences. Non-parametric Mann-Whitney U tests were employed for individual comparisons for datasets that significantly deviated from normality, as evidenced by strong departures from the expected diagonal line in Q-Q plots and significant Shapiro-Wilk test results (p < 0.05).

## Data availability

All RNA-sequencing and mass spectrometry data can be accessed through the NINDS gene GEO server (accession ID GSE281218) and the ProteomeXchange Consortium via the PRIDE partner repository (dataset identifier: PXD055701), respectively. Additional data can be obtained through contacting the authors upon reasonable request.

---

**Supplementary Figure 1.**
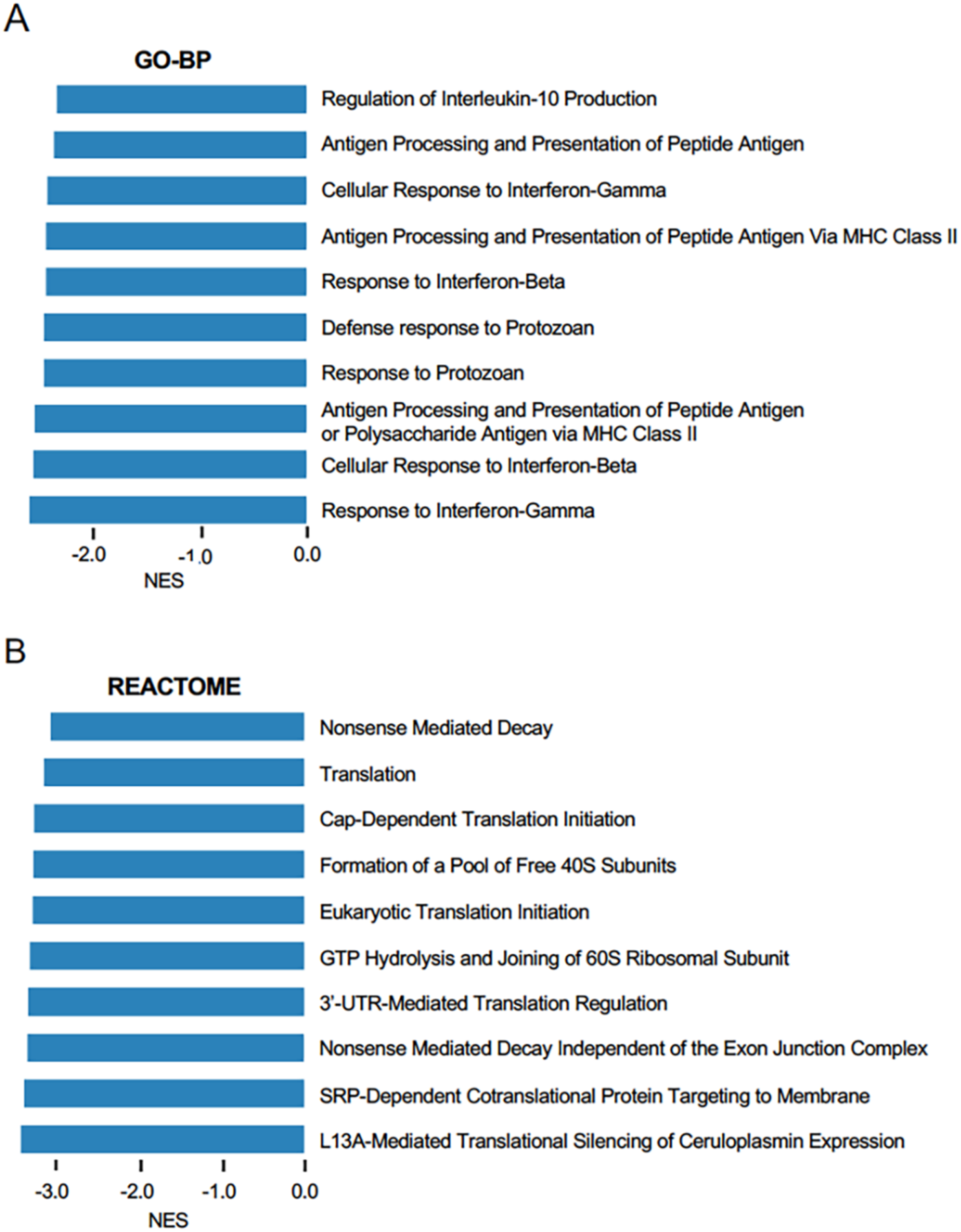
Downregulated pathways in P301S/TIA1cKO mice compared to P301S. A) Top 10 Gene Ontology-Biological Processes (GO-BP) pathways downregulated in P301S/TIA1cKO mice with normalized enrichment score (NES)>2. B) Top 10 Reactome pathways downregulated in P301S/TIA1cKO mice with normalized enrichment score >2.

**Supplementary Figure 2.**
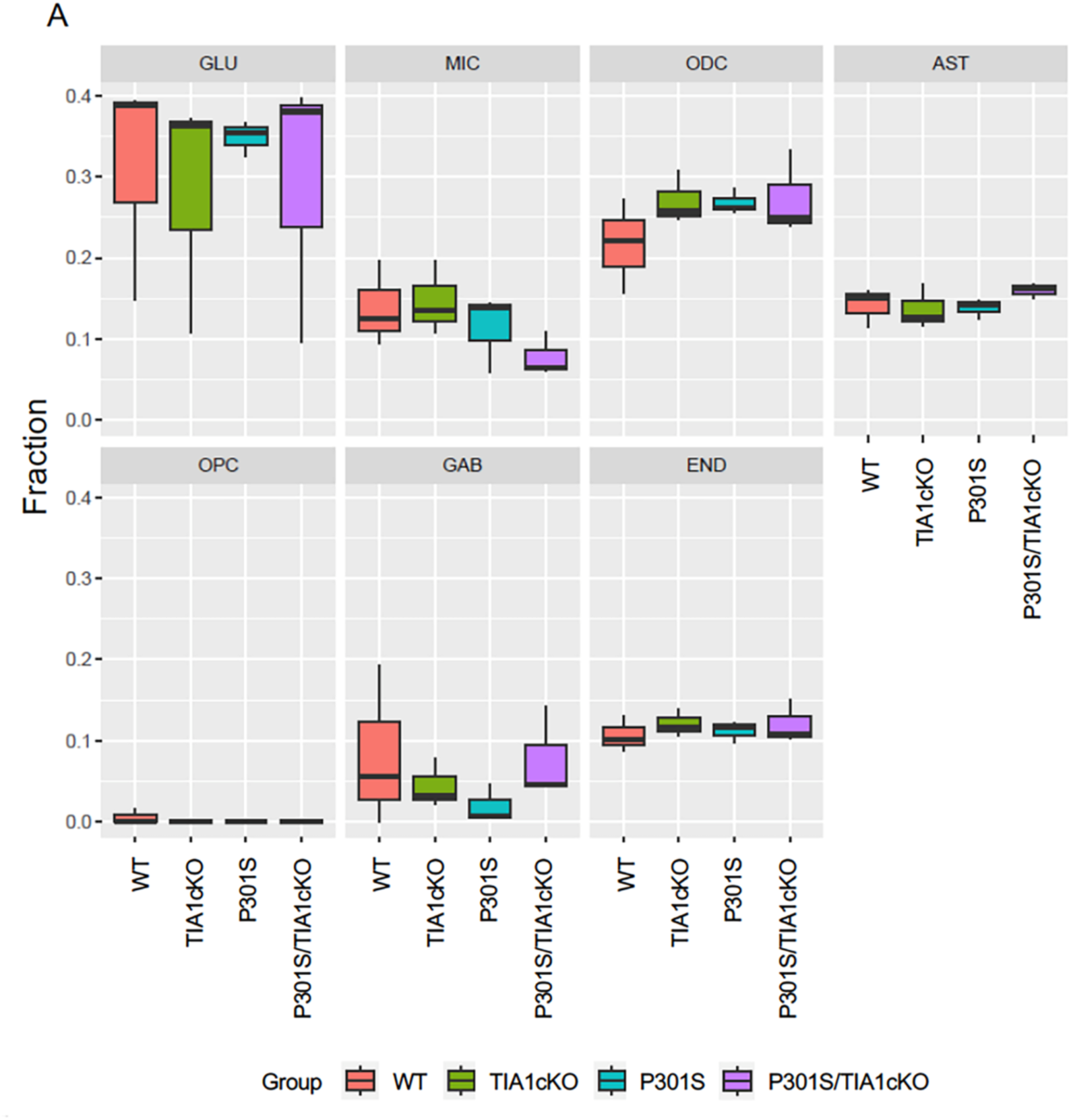
Microglial selective TIA1 knockout produces no differences in the proportions of specific cell-types. A) DeTREM deconvoluted data from WT, TIA1cKO, P301S, and P301S/TIA1cKO mice into proportion of glutamatergic neurons (GLU), microglia (MIC), oligodendrocytes (ODC), astrocytes (AST), oligodendrocytes precursors (OPC), GABAergic neurons (GAB), and endothelium (END).

**Supplementary Figure 3.**
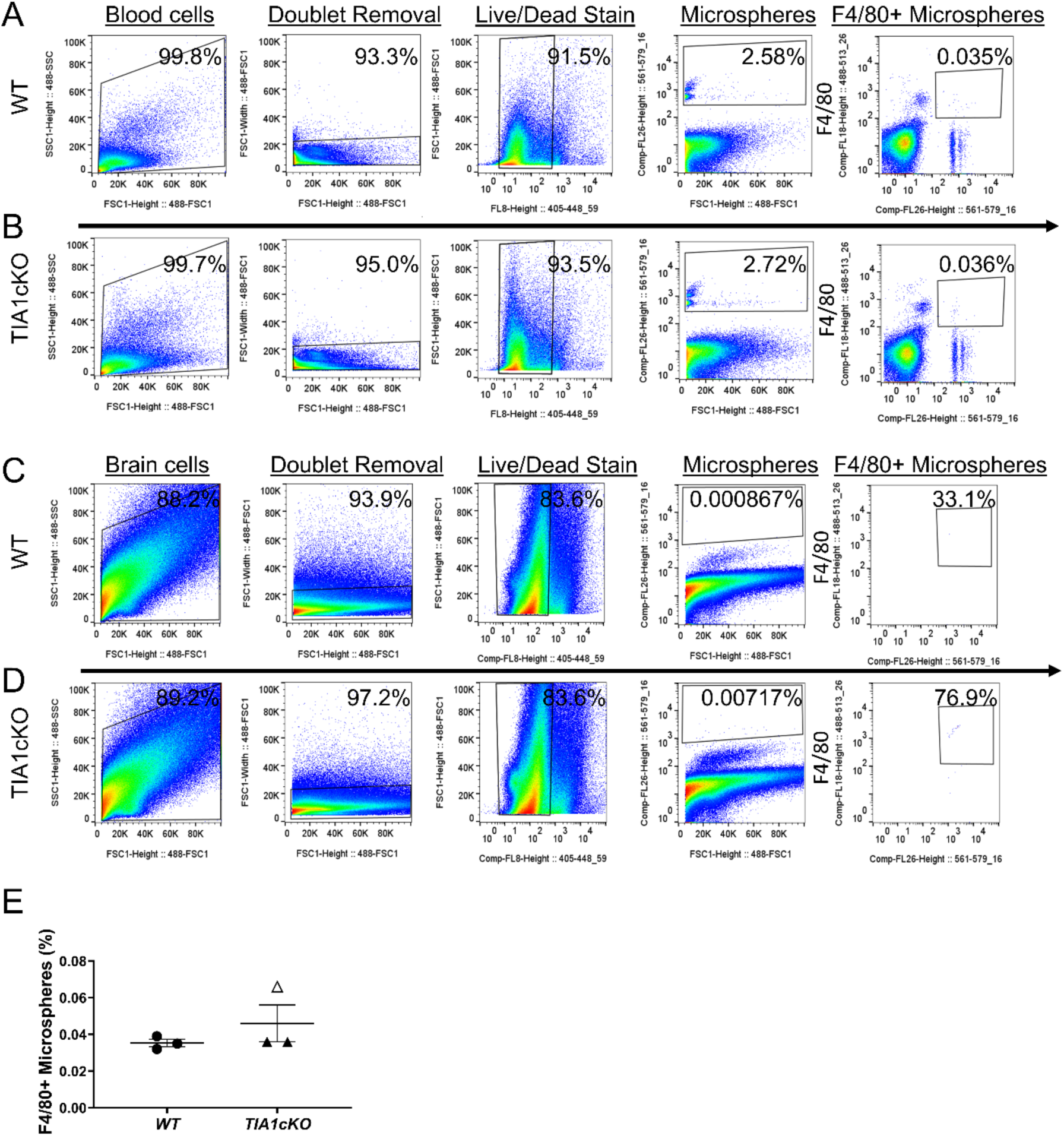
Microsphere uptake in blood versus brain. Fluorescent microspheres in the blood and brain 24 hours after intravenous injection and 3 hours post LPS injection. Gating strategy for microspheres blood of A) Wildtype (WT) and B) TIA1cKO mice. The proportion of microspheres that are found in F4/80+ monocytes in the blood of A) WT and B) TIA1cKO mice. Fluorescent microspheres found in brain of C) WT and D) TIA1cKO mice. The proportion of microspheres taken up by F4/80+ monocytes found in the brain of C) WT and D) TIA1cKO mice. E) Quantification of the proportion of F4/80+ microspheres in the blood. Data from TIA1cKO mouse with clear triangle was injected with microspheres but was not labelled with F4/80 antibody, the datapoint is an extrapolation of the TIA1cKO ratio of the percentage of injected microspheres to the percentage of F4/80+ microspheres.

**Supplementary Figure 4.**
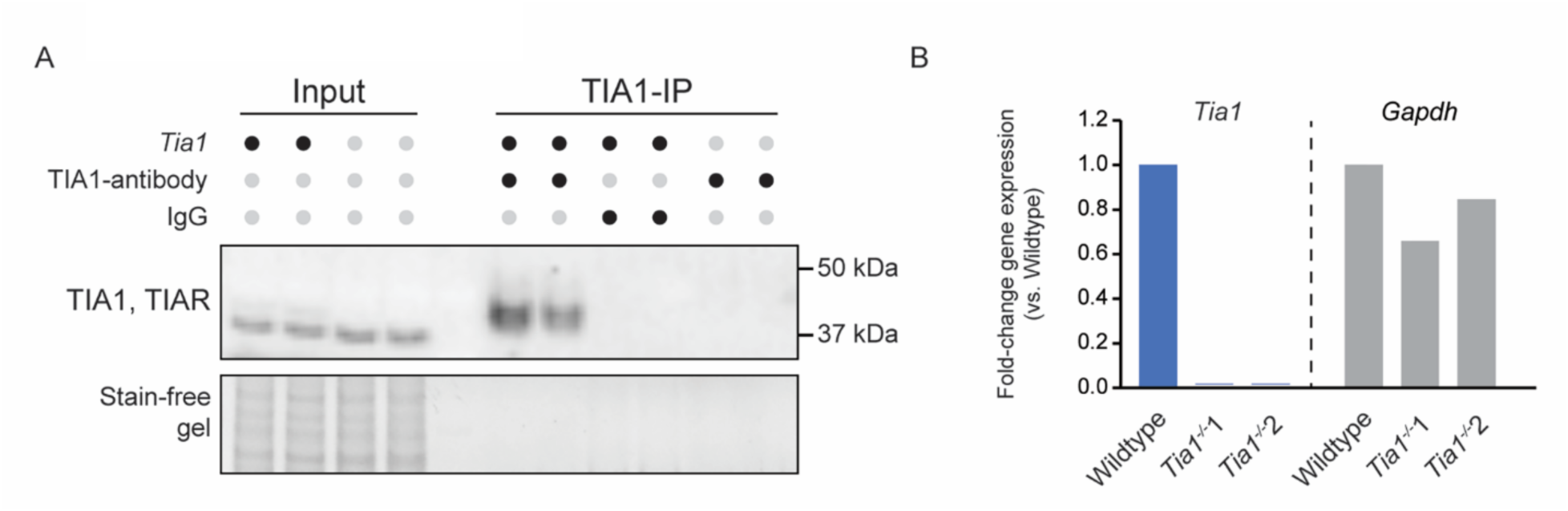
TIA1 antibody specificity testing for IP-MS. A) Immunopurification performed on wildtype and Tia1-/- brain lysates with TIA1-antibody (ab140595) and an IgG negative control. The blots stained with anti-TIA1/TIAR revealed successful pull-down of TIA1 in wildtypes only. B) qPCR on the Tia1-/- brain tissues revealing complete knock-out, indicating the immunoreactivity in Tia1-/- input lysates likely to be TIAR.

**Supplementary Figure 5.**
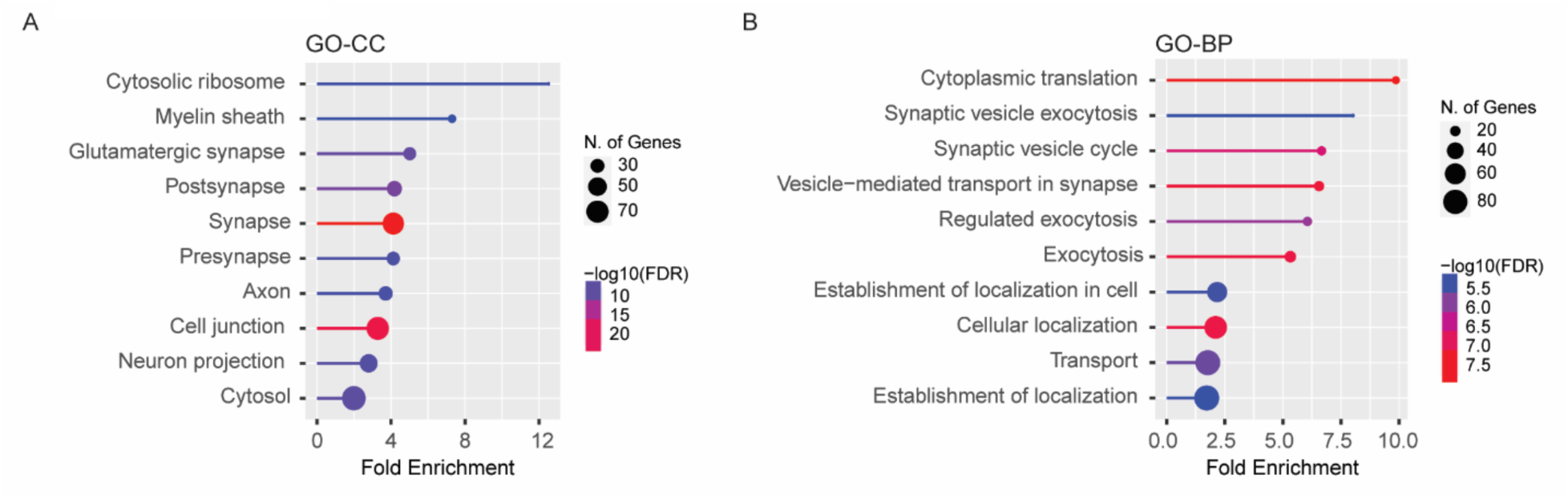
TIA1 interactome GO-enrichment analysis. A) TIA1-interactome shows enrichment for GO-terms related to ribosomal and synaptic processes as observed by the top-10 GO-Cellular Component terms, and B) GO-Biological Process terms.

**Supplementary Figure 6.**
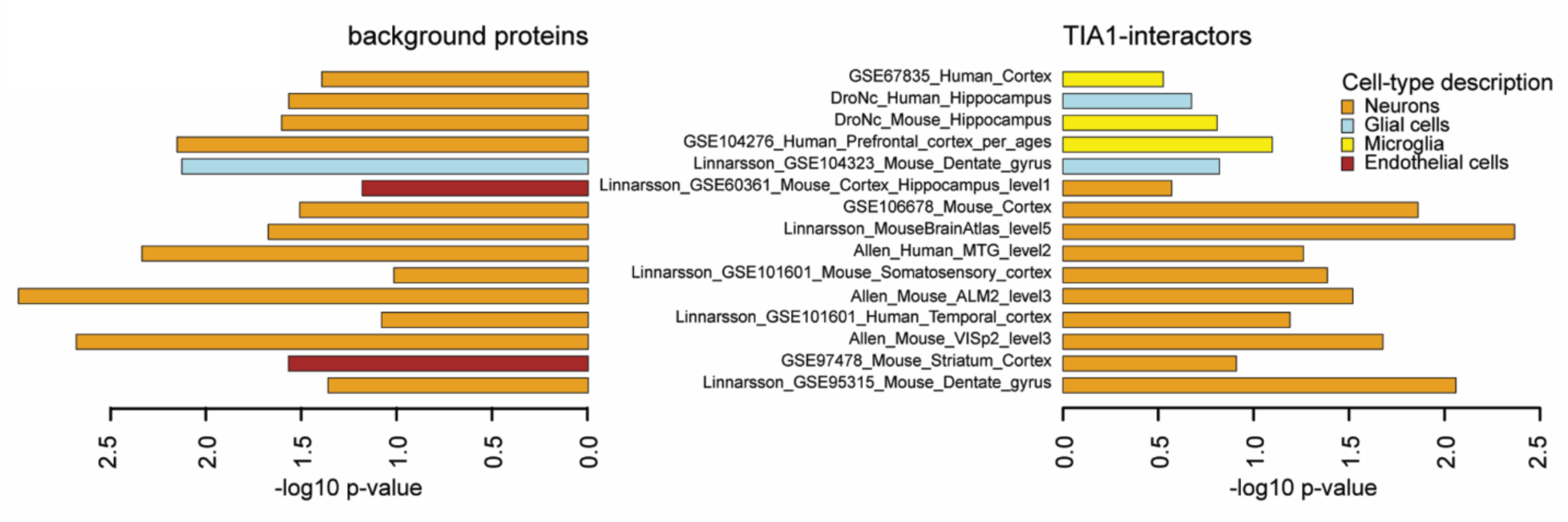
TIA1 interactome cell-type enrichment analysis. Expression weighted cell-type enrichment (EWCE) analysis using 15 previously published scRNAseq datasets from human and mouse cortical regions revealed neuronal and (micro)glial protein signatures in the TIA1-interactome. For each dataset the most statistically significant cell-type is shown. As a contrast, background proteins detected by IP-MS not considered true TIA1-interactors (e.g., due to high expression in the negative control IPs) did not reveal microglial protein signatures.

**Supplementary Figure 7.**
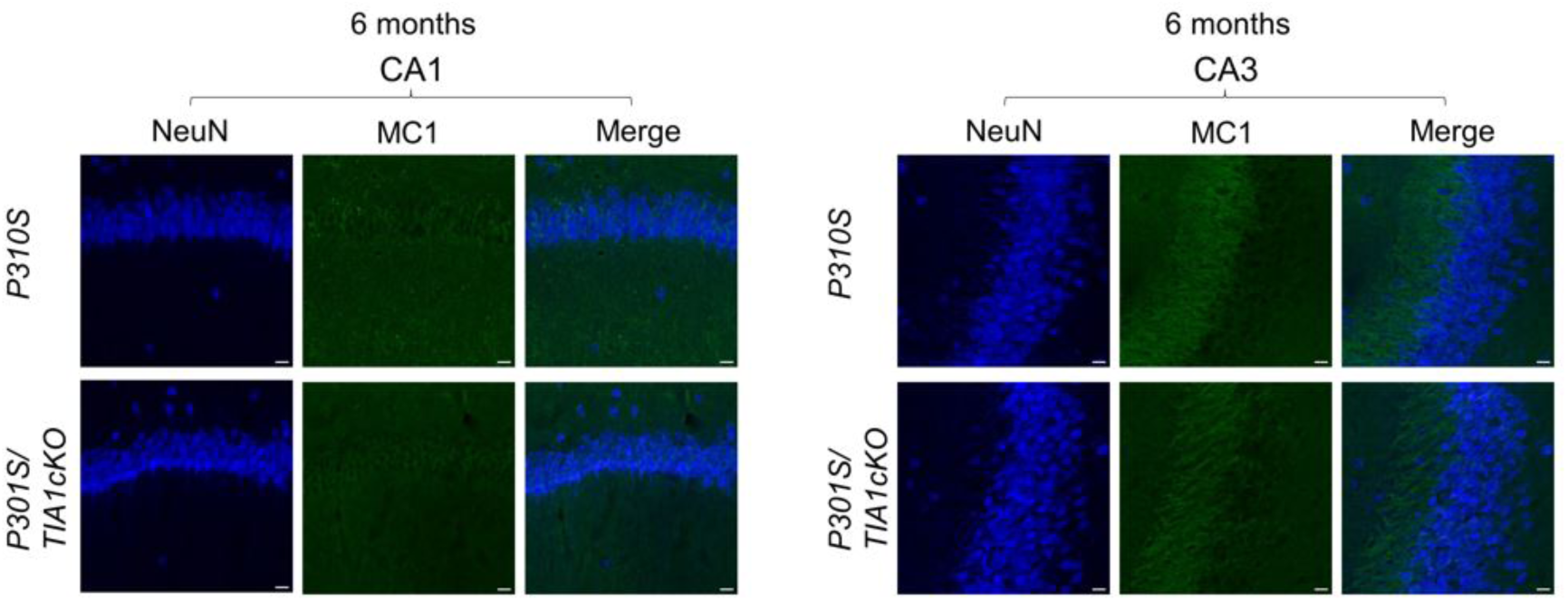
Tau pathology is minimal in 6-month-old P301S mice and not affected by microglial TIA1. A) CA1 region B) CA3 region of 6-month-old hippocampi from P301S and P301S/TIA1cKO mice stained for NeuN (405 nm) labelling neurons and MC1 (488 nm) labelling misfolded pathological tau. 20µm scale bars.

**Supplementary Figure 8.**
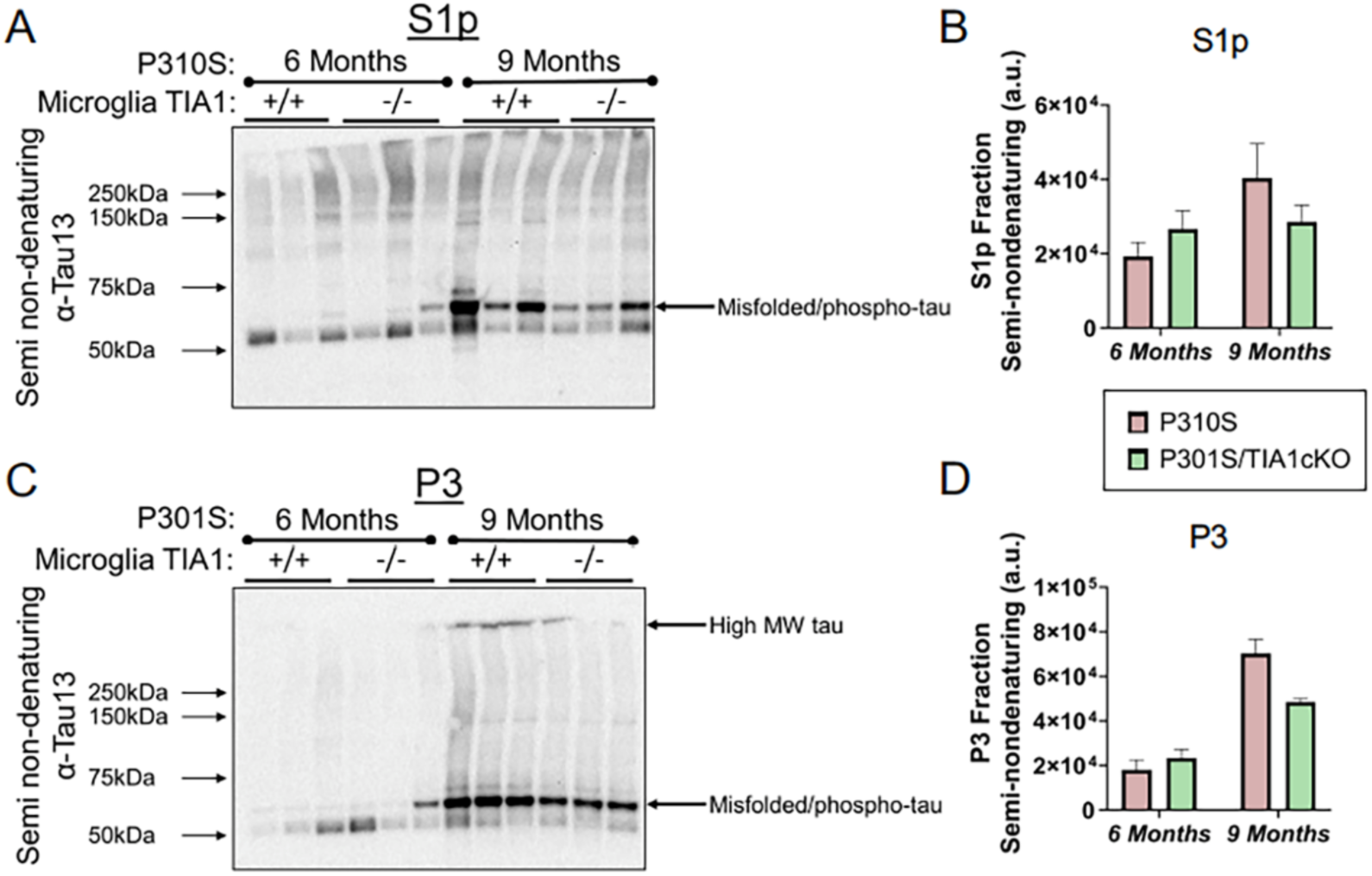
Fractionation of tau pathology from 6- and 9-month-old P301S and P301S/TIA1cKO mice. A) Semi-nondenaturing gel showing immunoblotting of total tau in the S1p fraction containing oligomeric tau using the antibody Tau13. The arrow identifies phosphorylated tau. B) Quantification of total tau in the S1p fraction. C) Semi-nondenaturing gel showing immunoblotting of total tau in the P3 fraction containing fibrillar tau using the antibody Tau13. The arrow identifies phosphorylated tau. D) Quantification of total tau in the P3 fraction.

## References

1. Rayman, J.B. & Kandel, E.R. TIA-1 Is a Functional Prion-Like Protein. Cold Spring Harb Perspect Biol 9(2017).

2. Meyer, C., et al. The TIA1 RNA-Binding Protein Family Regulates EIF2AK2-Mediated Stress Response and Cell Cycle Progression. Mol Cell 69, 622–635.e626 (2018).

3. Anderson, P., et al. A monoclonal antibody reactive with a 15-kDa cytoplasmic granule-associated protein defines a subpopulation of CD8+ T lymphocytes. J Immunol 144, 574–582 (1990).

4. Phillips, K., Kedersha, N., Shen, L., Blackshear, P.J. & Anderson, P. Arthritis suppressor genes TIA-1 and TTP dampen the expression of tumor necrosis factor alpha, cyclooxygenase 2, and inflammatory arthritis. Proc Natl Acad Sci U S A 101, 2011–2016 (2004).

5. Gilks, N., et al. Stress granule assembly is mediated by prion-like aggregation of TIA-1. Mol Biol Cell 15, 5383–5398 (2004).

6. Moreno, J.A., et al. Sustained translational repression by eIF2alpha-P mediates prion neurodegeneration. Nature 485, 507–511 (2012).

7. Wolozin, B. & Ivanov, P. Stress granules and neurodegeneration. Nat Rev Neurosci 20, 649–666 (2019).

8. Braak, H., Thal, D.R., Ghebremedhin, E. & Del Tredici, K. Stages of the pathologic process in Alzheimer disease: age categories from 1 to 100 years. J Neuropathol Exp Neurol 70, 960–969 (2011).

9. Perrin, R.J., Fagan, A.M. & Holtzman, D.M. Multimodal techniques for diagnosis and prognosis of Alzheimer’s disease. Nature 461, 916–922 (2009).

10. Therriault, J., et al. Biomarker modeling of Alzheimer’s disease using PET-based Braak staging. Nat Aging 2, 526–535 (2022).

11. Vanderweyde, T., et al. Contrasting Pathology of Stress Granule Proteins TIA-1 and G3BP in Tauopathies. J Neurosci 32, 8270–8283 (2012).

12. Vanderweyde, T., et al. Interaction of tau with the RNA-binding Protein TIA1 Regulates tau Pathophysiology and Toxicity. Cell Rep 15, 1–12 (2016).

13. Ash, P.E.A., et al. TIA1 potentiates tau phase separation and promotes generation of toxic oligomeric tau. Proc Natl Acad Sci U S A 118(2021).

14. Apicco, D.J., et al. Reducing the RNA binding protein TIA1 protects against tau-mediated neurodegeneration in vivo. Nat Neurosci 21, 72–80 (2018).

15. Li, Q. & Barres, B.A. Microglia and macrophages in brain homeostasis and disease. Nat Rev Immunol 18, 225–242 (2018).

16. Rodríguez, J.J., et al. Increased densities of resting and activated microglia in the dentate gyrus follow senile plaque formation in the CA1 subfield of the hippocampus in the triple transgenic model of Alzheimer’s disease. Neurosci Lett 552, 129–134 (2013).

17. Sheng, J.G., Mrak, R.E. & Griffin, W.S. Neuritic plaque evolution in Alzheimer’s disease is accompanied by transition of activated microglia from primed to enlarged to phagocytic forms. Acta Neuropathol 94, 1–5 (1997).

18. Paolicelli, R.C., et al. Microglia states and nomenclature: A field at its crossroads. Neuron 110, 3458–3483 (2022).

19. Hickman, S.E., et al. The microglial sensome revealed by direct RNA sequencing. Nat Neurosci 16, 1896–1905 (2013).

20. Hickman, S.E. & El Khoury, J. Analysis of the Microglial Sensome. Methods Mol Biol 2034, 305–323 (2019).

21. Akhter, R., Shao, Y., Formica, S., Khrestian, M. & Bekris, L.M. TREM2 alters the phagocytic, apoptotic and inflammatory response to Aβ(42) in HMC3 cells. Mol Immunol 131, 171–179 (2021).

22. Krance, S.H., et al. The complement cascade in Alzheimer’s disease: a systematic review and meta-analysis. Mol Psychiatry (2019).

23. Asai, H., et al. Depletion of microglia and inhibition of exosome synthesis halt tau propagation. Nat Neurosci 18, 1584–1593 (2015).

24. Spangenberg, E.E. & Green, K.N. Inflammation in Alzheimer’s disease: Lessons learned from microglia-depletion models. Brain Behav Immun 61, 1–11 (2017).

25. Keren-Shaul, H., et al. A Unique Microglia Type Associated with Restricting Development of Alzheimer’s Disease. Cell 169, 1276–1290 e1217 (2017).

26. Krasemann, S., et al. The TREM2-APOE Pathway Drives the Transcriptional Phenotype of Dysfunctional Microglia in Neurodegenerative Diseases. Immunity 47, 566–581.e569 (2017).

27. Dallas, M.L. & Widera, D. TLR2 and TLR4-mediated inflammation in Alzheimer’s disease: self-defense or sabotage? Neural Regen Res 16, 1552–1553 (2021).

28. Hong, S., et al. Complement and microglia mediate early synapse loss in Alzheimer mouse models. Science 352, 712–716 (2016).

29. Stephan, A.H., Barres, B.A. & Stevens, B. The complement system: an unexpected role in synaptic pruning during development and disease. Annu Rev Neurosci 35, 369–389 (2012).

30. Zhou, J., Fonseca, M.I., Pisalyaput, K. & Tenner, A.J. Complement C3 and C4 expression in C1q sufficient and deficient mouse models of Alzheimer’s disease. J Neurochem 106, 2080–2092 (2008).

31. Gomez-Arboledas, A., et al. C5aR1 antagonism alters microglial polarization and mitigates disease progression in a mouse model of Alzheimer’s disease. Acta Neuropathol Commun 10, 116 (2022).

32. O’Neill, N.K., et al. Bulk brain tissue cell-type deconvolution with bias correction for single-nuclei RNA sequencing data using DeTREM. BMC Bioinformatics 24, 349 (2023).

33. Grubman, A., et al. Transcriptional signature in microglia associated with Aβ plaque phagocytosis. Nat Commun 12, 3015 (2021).

34. Ma, W., Huang, G., Wang, Z., Wang, L. & Gao, Q. IRF7: role and regulation in immunity and autoimmunity. Front Immunol 14, 1236923 (2023).

35. Blank, T. & Prinz, M. Type I interferon pathway in CNS homeostasis and neurological disorders. Glia 65, 1397–1406 (2017).

36. Masood, K.I., et al. Upregulated type I interferon responses in asymptomatic COVID-19 infection are associated with improved clinical outcome. Sci Rep 11, 22958 (2021).

37. De Rossi, P., et al. Predominant expression of Alzheimer’s disease-associated BIN1 in mature oligodendrocytes and localization to white matter tracts. Mol Neurodegener 11, 59 (2016).

38. De Rossi, P., et al. Neuronal BIN1 Regulates Presynaptic Neurotransmitter Release and Memory Consolidation. Cell Rep 30, 3520–3535.e3527 (2020).

39. Saha, O., et al. The Alzheimer’s disease risk gene BIN1 regulates activity-dependent gene expression in human-induced glutamatergic neurons. Mol Psychiatry 29, 2634–2646 (2024).

40. Li, X., Rayman, J.B., Kandel, E.R. & Derkatch, I.L. Functional role of tia1/pub1 and sup35 prion domains: directing protein synthesis machinery to the tubulin cytoskeleton. Mol Cell 55, 305–318 (2014).

41. Rayman, J.B., et al. Genetic Perturbation of TIA1 Reveals a Physiological Role in Fear Memory. Cell Rep 26, 2970–2983 e2974 (2019).

42. DeVries, S.A., et al. Immune proteins C1q and CD47 may contribute to aberrant microglia-mediated synapse loss in the aging monkey brain that is associated with cognitive impairment. Geroscience 46, 2503–2519 (2024).

43. Werneburg, S., et al. Targeted Complement Inhibition at Synapses Prevents Microglial Synaptic Engulfment and Synapse Loss in Demyelinating Disease. Immunity 52, 167–182 e167 (2020).

44. Miron, J., Picard, C., Labonté, A., Auld, D. & Poirier, J. MSR1 and NEP Are Correlated with Alzheimer’s Disease Amyloid Pathology and Apolipoprotein Alterations. J Alzheimers Dis 86, 283–296 (2022).

45. Rosenzweig, N., et al. PD-1/PD-L1 checkpoint blockade harnesses monocyte-derived macrophages to combat cognitive impairment in a tauopathy mouse model. Nat Commun 10, 465 (2019).

46. Scott-Hewitt, N., et al. Microglial-derived C1q integrates into neuronal ribonucleoprotein complexes and impacts protein homeostasis in the aging brain. Cell 187, 4193–4212.e4124 (2024).

47. Kim, H.S., et al. Distinct binding properties of TIAR RRMs and linker region. RNA Biol 10, 579–589 (2013).

48. Suswam, E.A., Li, Y.Y., Mahtani, H. & King, P.H. Novel DNA-binding properties of the RNA-binding protein TIAR. Nucleic Acids Res 33, 4507–4518 (2005).

49. Kishore, U. & Reid, K.B. C1q: structure, function, and receptors. Immunopharmacology 49, 159–170 (2000).

50. Zhuang, B., Mancarci, B.O., Toker, L. & Pavlidis, P. Mega-Analysis of Gene Expression in Mouse Models of Alzheimer’s Disease. eNeuro 6(2019).

51. Liddelow, S.A., et al. Neurotoxic reactive astrocytes are induced by activated microglia. Nature 541, 481–487 (2017).

52. Van Hove, H., et al. A single-cell atlas of mouse brain macrophages reveals unique transcriptional identities shaped by ontogeny and tissue environment. Nat Neurosci 22, 1021–1035 (2019).

53. Lu, Y.C., Yeh, W.C. & Ohashi, P.S. LPS/TLR4 signal transduction pathway. Cytokine 42, 145–151 (2008).

54. Lysakova-Devine, T., et al. Viral inhibitory peptide of TLR4, a peptide derived from vaccinia protein A46, specifically inhibits TLR4 by directly targeting MyD88 adaptor-like and TRIF-related adaptor molecule. J Immunol 185, 4261–4271 (2010).

55. Lee, A.J., Ro, M., Cho, K.J. & Kim, J.H. Lipopolysaccharide/TLR4 Stimulates IL-13 Production through a MyD88-BLT2-Linked Cascade in Mast Cells, Potentially Contributing to the Allergic Response. J Immunol 199, 409–417 (2017).

56. Shi, L., et al. IL-1 Transcriptional Responses to Lipopolysaccharides Are Regulated by a Complex of RNA Binding Proteins. J Immunol 204, 1334–1344 (2020).

57. Schoggins, J.W., et al. Corrigendum: A diverse range of gene products are effectors of the type I interferon antiviral response. Nature 525, 144 (2015).

58. Kang, Q., Li, L., Pang, Y., Zhu, W. & Meng, L. An update on Ym1 and its immunoregulatory role in diseases. Front Immunol 13, 891220 (2022).

59. Aerbajinai, W., Lee, K., Chin, K. & Rodgers, G.P. Glia maturation factor-gamma negatively modulates TLR4 signaling by facilitating TLR4 endocytic trafficking in macrophages. J Immunol 190, 6093–6103 (2013).

60. Lotz, M., et al. Amyloid beta peptide 1-40 enhances the action of Toll-like receptor-2 and −4 agonists but antagonizes Toll-like receptor-9-induced inflammation in primary mouse microglial cell cultures. J Neurochem 94, 289–298 (2005).

61. Zhu, B., et al. Trem2 deletion enhances tau dispersion and pathology through microglia exosomes. Mol Neurodegener 17, 58 (2022).

62. Young, K. & Morrison, H. Quantifying Microglia Morphology from Photomicrographs of Immunohistochemistry Prepared Tissue Using ImageJ. J Vis Exp (2018).

63. Campos, A.B., Duarte-Silva, S., Ambrósio, A.F., Maciel, P. & Fernandes, B.

64. Dobin, A., et al. STAR: ultrafast universal RNA-seq aligner. Bioinformatics 29, 15–21 (2013).

65. Liao, Y., Smyth, G.K. & Shi, W. The Subread aligner: fast, accurate and scalable read mapping by seed-and-vote. Nucleic Acids Res 41, e108 (2013).

66. Love, M.I., Huber, W. & Anders, S. Moderated estimation of fold change and dispersion for RNA-seq data with DESeq2. Genome Biol 15, 550 (2014).

67. Subramanian, A., et al. Gene set enrichment analysis: a knowledge-based approach for interpreting genome-wide expression profiles. Proc Natl Acad Sci U S A 102, 15545–15550 (2005).

68. Merico, D., Isserlin, R., Stueker, O., Emili, A. & Bader, G.D. Enrichment map: a network-based method for gene-set enrichment visualization and interpretation. PLoS One 5, e13984 (2010).

69. Lau, S.F., Cao, H., Fu, A.K.Y. & Ip, N.Y. Single-nucleus transcriptome analysis reveals dysregulation of angiogenic endothelial cells and neuroprotective glia in Alzheimer’s disease. Proc Natl Acad Sci U S A 117, 25800–25809 (2020).

70. Blake, J.A., et al. Mouse Genome Database (MGD): Knowledgebase for mouse-human comparative biology. Nucleic Acids Res 49, D981–d987 (2021).

71. Chen, Z., et al. seq-ImmuCC: Cell-Centric View of Tissue Transcriptome Measuring Cellular Compositions of Immune Microenvironment From Mouse RNA-Seq Data. Front Immunol 9, 1286 (2018).

72. Newman, A.M., et al. Determining cell type abundance and expression from bulk tissues with digital cytometry. Nat Biotechnol 37, 773–782 (2019).

73. Ge, S.X., Jung, D. & Yao, R. ShinyGO: a graphical gene-set enrichment tool for animals and plants. Bioinformatics 36, 2628–2629 (2020).

74. Watanabe, K., Umićević Mirkov, M., de Leeuw, C.A., van den Heuvel, M.P. & Posthuma, D. Genetic mapping of cell type specificity for complex traits. Nat Commun 10, 3222 (2019).

